# Modeling myosin with interacting linkages

**DOI:** 10.1101/2021.04.20.440673

**Authors:** Tosan Omabegho

## Abstract

In this study, I describe a model in which mechanical linkages dynamically interact in a stepwise and reversible manner, and use it to model the chemical cycle and lever arm action of the biomolecular motor myosin. Myosin is emulated using a series of multivalent chemical reactions between a linkage enzyme and four reactants: a cleaveable fuel, two cleavage products, and ligand. Geometric coupling between the fuel and ligand binding sites—an analog for negative allosteric coupling—allows reaction sequences similar to nucleotide exchange to take place that in turn drive the “strokes” of the machine’s lever arm. Cyclic chemical behavior is demonstrated by stochastic simulation, and mechanical activity by a series of logical arguments. I show how a reciprocal and nonreciprocal conformational cycle emerge from the allosteric rules designed to achieve chemical cycling, and how the non-reciprocal cycle can break directional symmetry along a track like structure. A dimeric construct is used to demonstrate how directed motion can be designed by inhibition of the reciprocal cycle and reinforcement of the non-reciprocal cycle, through allosteric feedback between the units of the dimer. By showing how the chemomechanical cycle of a biomolecular motor can be recreated with simple geometric and chemical principles, this work may help advance the rational design of allosteric mechanisms, and the development of synthetic molecular motors.

## 1 Introduction

Biomolecular motors are autonomous nanoscale machines that use chemical energy to produce directed motion [1]. While much has been learned about biomolecular motors, fundamental questions remain about how they operate. Fundamental questions remain is the sense that it is still not clear how to construct similarly operating synthetic motors [2–4].

One key mechanism that is not well understood is allostery [5, 6]. Allostery is defined as intramolecular communication between binding sites [6–8]. The mechanism allows molecular motors, each of which accomplishes different mechanical tasks, to use the same chemical (ATP) as fuel. Although allostery is intramolecular communication, intermolecular reactions often begin and end allosteric reaction sequences, allowing information to flow through molecular motors. This ‘flow-through’ produces the chemical and conformational changes that accompany sequential binding reactions. However, there are no models that capture the intramolecular and intermolecular aspects of allosteric reactions in molecular motors, which limits the extent to which we can emulate nature.

To address this gap, I recreate the chemical and mechanical action of the motor myosin using a model in which mechanical linkages ‘chemically’ interact through multivalent contacts (‘chemo-mechanical linkages’). Motivated by an earlier version of this paper, reference [9] developed a formal model of chemo-mechanical linkages. They also showed that complex behavior can be achieved in an idealized topological model, which considers solely the graph connectivity of the linkages.

Myosins use ATP to either generate directed motion along actin filament tracks, or move actin filaments relative to a fixed reference frame, as they do in muscle [10]. Their directed motion depends upon the movement of a rigid protrusion called the lever arm, which amplifies conformational changes that originate in the main body (or head) of the enzyme [11], which contains the ATP and actin binding domains. The conformational changes that produce myosin’s lever arm motion take place during nucleotide exchange, when either ATP is loaded onto the motor, or the waste products of hydrolysis (Pi and ADP) are expelled from the motor [10, 12, 13]. Each stage is controlled by allosteric communication between myosin’s ATP and actin binding sites.

In short, when ATP is loaded onto a motor unit, it causes the head to detach from actin and the recovery stroke, in which the orientation of the lever arm is reset. And conversely, when a head binds actin, it causes the head to expel Pi and ADP, and the power stroke, in which the orientation of the lever arm is set in the forward direction [12]. This tight coupling, between nucleotide exchange (chemical events) and lever arm motion (mechanical events), suggests that reproducing the chemo-mechanical action of myosin requires reproducing the allosteric communication that enables nucleotide exchange.

Following this line of reasoning, I represent myosin and its four reactants (ATP, Pi, ADP, and actin) as a set of interacting linkages, and construct a mechanism in which competing multivalent binding reactions to the myosin linkage, emulate negative allosteric coupling between two binding sites, and drive conformational change. (The idea that multivalent binding drives conformational change in a molecular motor is not new [14, 15].) The negative coupling mechanism allows reaction sequences, referred to here as *allosteric displacements* [16], to take place that are analogs for nucleotide exchange. I describe two allosteric displacement sequences, one controlling for fuel loading and one product exhaust. The two sequences are connected together by a simple model of catalysis, enabling the system to autonomously cycle and emulate the chemical cycle of a myosin monomer.

By adding a lever arm to the enzyme linkage, I show how the catalytic cycle is able to drive reciprocal and non-reciprocal lever arm motion on a track-like structure. As in myosin, the machine’s recovery stroke takes place during fuel loading, and its power stroke during product exhaust. However, in contrast to myosin, its recovery stroke takes place before it dissociates from its actin analog, through a conformational change of the enzyme head relative to a fixed lever arm position. Finally, I describe how allosteric feedback between units in a dimeric construct can bias track-detached units to bind in the forward position, which is a mechanism for generating directed motion along a track [17, 18]. The feedback results in selection of the non-reciprocal conformational cycle over the reciprocal cycle.

This article is organized as follows: First, I describe the components and operation of the linkage system, and stochastic simulations that demonstrate how the linkage system emulates the chemical cycle of a myosin monomer. Futile “off-target” behaviors are also analyzed. Second, I describe how reciprocal and non-reciprocal lever arm cycles can be mapped onto the system. Lastly, I describe the design of the dimer construct that is a directed walker.

## Results

### Operation of the linkage system

The myosin monomer system comprises five molecules (myosin, ATP, Pi, ADP and actin) that interact in a sequence of six chemical steps, which are emulated here with mechanical linkages. These six steps can be grouped into two nucleotide exchange sequences (or displacement sequences), named displacement 1 and displacement 2, which are separated by the hydrolysis reaction:

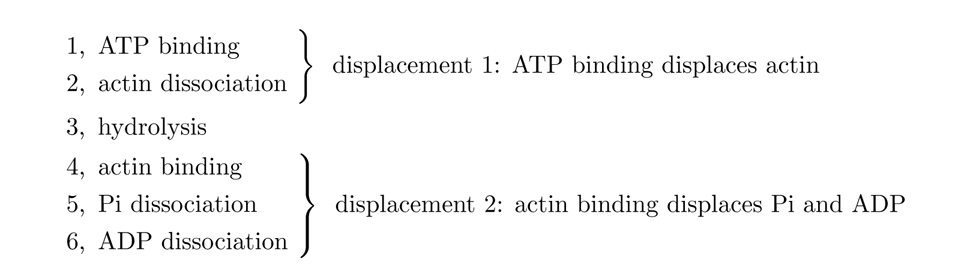

The five molecules of the linkage system (Fig. 1A) map to the myosin molecules like so:

**Fig. 1.**
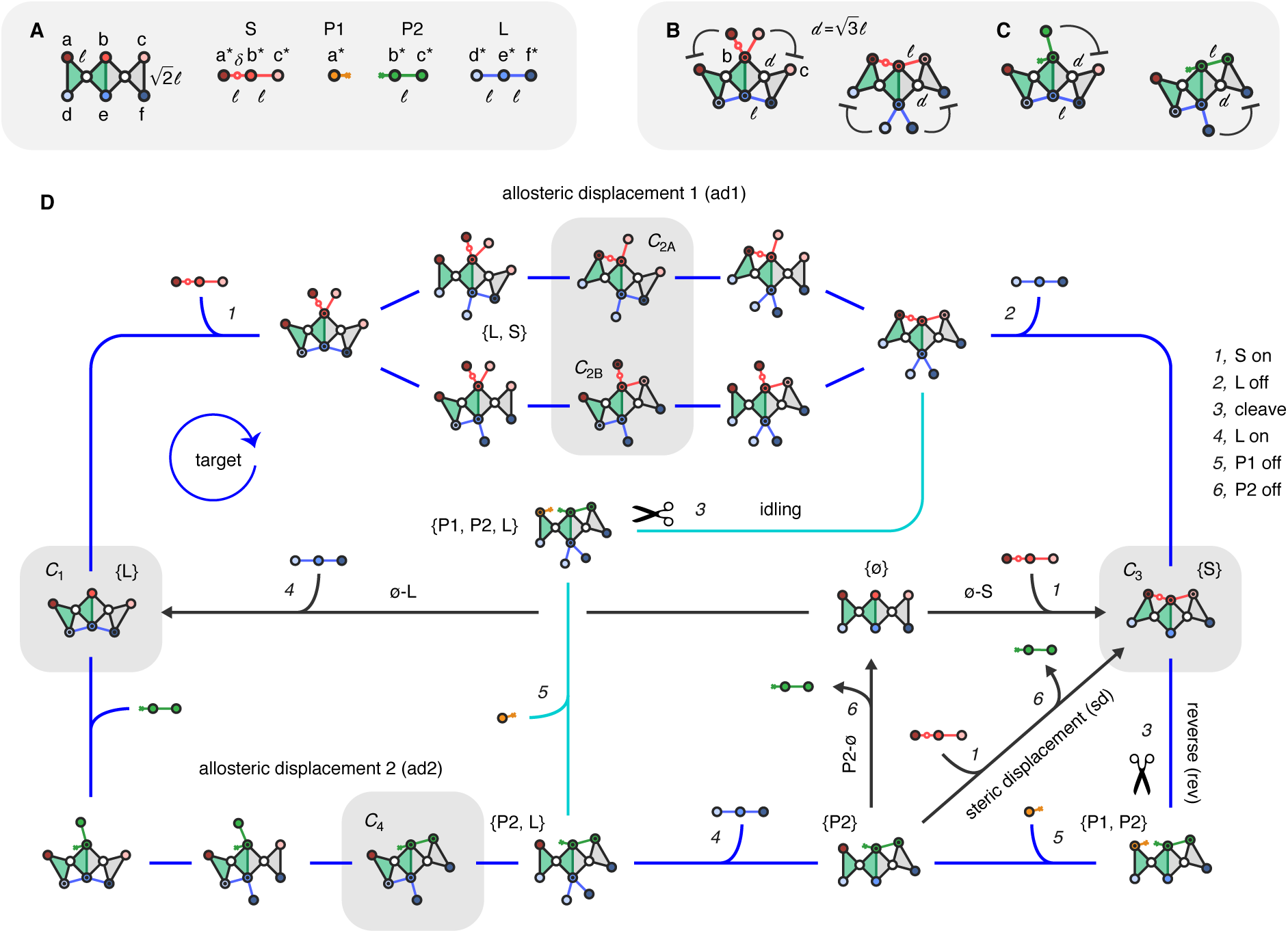
The linkage system. **A,** Five molecules of the linkage system: the enzyme, S, P1, P2 and L. The enzyme is a chain of two linkage unit (one green, one gray) fused together on edge, where each unit consists of two right triangles connected at a rotatable node (white). Edge lengths are *ℓ* and 2 *ℓ*. Three red vertices (a, b and c; top) form the binding site for S, P1 and P2; and three blue vertices (d, e and f; bottom) form the binding site for L. The left side (green) is catalytic, and cuts S into P1 and P2, while the right side (gray) is non-catalytic. S contains three complimentary nodes (a*, b* and c*) to the enzyme separated by two bars of length *ℓ*. S is cut by the enzyme at the special node *δ*. P1 contains node a*, and P2 contains nodes a* and b*. L contains nodes d*, e* and f*. **B,** Left, L blocks S. L blocks from binding by bending the enzyme downwards when bound trivalently, placing S’s nodes out of reach (blunt arrowheads). L, S and P2 all contain bars of length *ℓ*, and thus spread the nodes at the opposing binding site to length *d* = 3 *ℓ*. Right, S blocks L. **C,** Left, L blocks P2. Right, P2 blocks L. **D,** Target cycle and futile pathways. The target cycle (outside blue path; forward is clockwise) takes place in thirteen reversible transitions, six of which are intermolecular (numbered) and map to myosin (see main text). Starting at L, the allosteric displacement of L by S (‘ad1’) takes place along two pathways in four intramolecular steps in the interval between 1 and 2 ([1, 2]), leaving the enzyme bound to S (S). Following cleavage (3), P1 dissociates randomly (5), leaving the enzyme bound to P2. The displacement of P2 by L (‘ad2’) takes place in three intramolecular steps between 4 and 6 ([1, 2]), returning the system to state L. See the main text for a description of the futile pathways (idling, P2-ø, ø-L, ø-S, and steric displacement). The five states highlighted in gray and specifically named (*C*_1_*, C*_2*A*_*, C*_2*B*_*, C*_3_ and *C*_4_) are rigid. Arrow notation: headless arrows denote single, reversible transitions (all steps along target and idling); headed arrows represent two or more transitions between states (along P2-ø, ø-L, ø-S, and steric displacement).

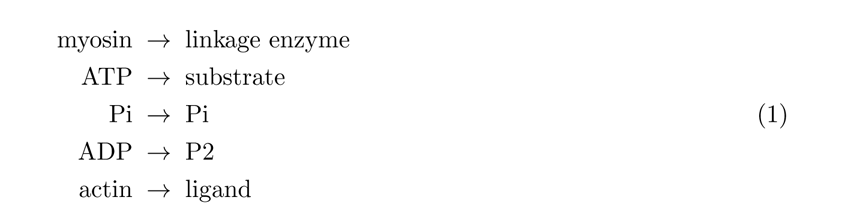

The linkage enzyme is a chain of two domains *√*(o ne green, one gray) fused together on their long edges. The short edges are length *ℓ* and the long edges 2 *ℓ*. Each domain consists of two isosceles right triangles connected at a rotatable node (white). This node connection gives each domain one degree of freedom, and the two-domain enzyme two degrees of freedom. The six remaining nodes, three on top and three on the bottom, form multivalent binding sites on top for the substrate, P1 and P2, and on the bottom for the ligand. Each node on the enzyme makes a chemically specific and reversible bond with a complimentary node on a reactant. Complimentarity is defined by unstarred and starred letter pairs on the enzyme and reactants, respectively. The node pairings allow substrate to bind trivalently at a, b and c, P1 monovalently at a, P2 divalently at b and c, and ligand trivalently d, e and f. Substrate contains an additional special node *δ*, between *a\** and *b\**, where it is either cleaved by the enzyme into P1 and P2, or fused together by the enzyme to form substrate from P1 and P2.

Negative allosteric coupling works like this. Because substrate, P2 and ligand are each composed of nodes connected by bars of length *ℓ*, they bend the enzyme in opposing conformatio*√*ns (Fig. 1B and C). This bending increases the distance between nodes on the opposite side to length *d* = 3 *ℓ*, such that if a molecule is bound to any single node on the opposite side, it cannot reach a second node. In this way, substrate and ligand are negatively coupled on both the left and right side of the enzyme (Fig. 1B), and P2 and the ligand are negatively coupled on the right side only—because P2 only binds divalently on the right side (Fig. 1C). P1, as a single node, cannot affect the geometry of the enzyme and so it not allosterically active. These relationships can be summarized in a table form (myosin is included for comparison on the right):

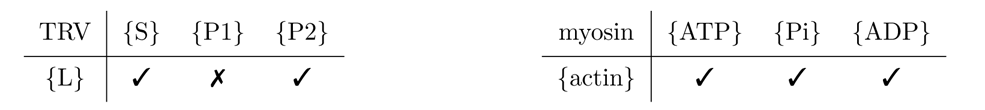

The check-marks mean the molecules are negatively coupled, and ‘x’ means they are not. As indicated, the main difference between allosteric coupling in the linkage and myosin systems is that P1 (the Pi analog) is not negatively coupled to ligand (the actin analog), whereas Pi in myosin is negatively coupled to actin.

The allosteric coupling and myosin-to-linkage species mapping defined in eq. 1 can be used to map the linkage system to the six myosin intermolecular reactions like so:

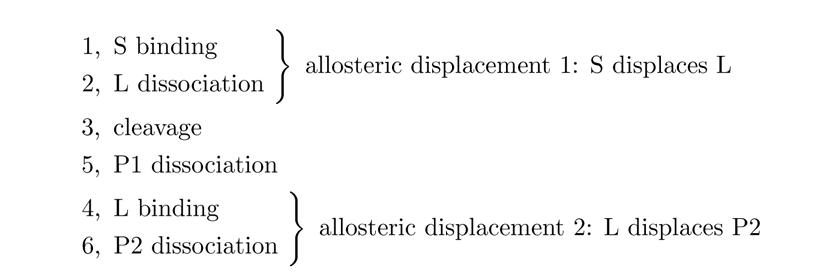

This sequence is the target intermolecular behavior of the system. Note that because P1 binds to the enzyme at a single node, its dissociation (reaction 5) comes before ligand binding (reaction 4). But, this rule is not a strict rule. What this ordering means, is that P1’s dissociation is not coupled ligand binding—P1 dissociates randomly—and therefore reaction 5 can take place anytime after P1 is created by the cleavage reaction (reaction 3).

The complete target path (Fig. 1D, blue), which includes intermolecular and intramolecular transitions, is a completely reversible sequence (hence, no arrow heads) of discrete transitions. The forward direction is clockwise. The transition rule followed along the path is that transitions between states can be made if two sates differ by one more, or one less node connection. Starting at state L, substrate binds and displaces ligand in a sequence of four steps, along each of the two mechanical path that can be defined by the symmetry of the reaction (see SI 2.1 for all paths). Steps alternate between ligand dissociation and substrate binding. By the end of the sequence, substrate binding has converted the ligand association to a single node interaction, and overtaken the two mechanical degrees of freedom. After the cleavage reaction and P1 dissociation, ligand rebinds and displaces P2 in a three-step sequence. A divalent connection is made in the first step (by contrast to substrate displacing ligand), followed by alternating displacement steps, in which P2 dissociates, and ligand binds trivalently to overtake the right mechanical degree of freedom. This final step returns the system to state *{*L*}*. [The five states highlighted in gray and given specific names (*C*_1_*, C*_2*A*_*, C*_2*B*_*, C*_3_ and *C*_4_) are completely rigid states that occur along the target cycle. These states will be used later to define the mechanical behavior of the system.]

Forward cycling along the target path is achieved by assigning transition rates according to three guidelines, which are described in the next section. Diversions from the target path, along which substrate is not constructively used, are completed along five main futile events (Fig. 1D, pathways drawn on the inside). These events are:

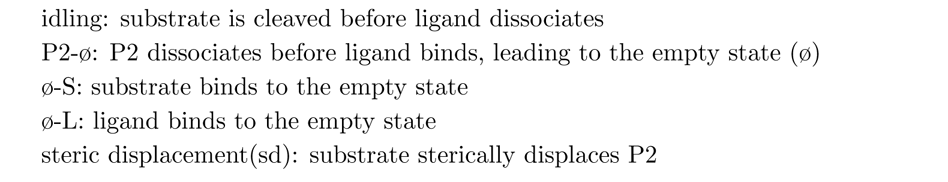

Except for idling, arrowheads are used to indicate that these futile events are depicted without their intervening steps (see SI 2.2 for complete paths). Each futile event ruins at least one allosteric displacement sequence, and thus one stage of nucleotide exchange, which becomes uncoupled from its corresponding ligand exchange process. Along idling, substrate loads but fails to displace ligand. This event means that the same ligand molecule present at the start of a fuel cycle is present at the end. Thus, the first stage of ligand exchange, which is ligand dissociation, does not take place as a result of idling. Along the other four futile events, P2 dissociates before ligand binds, and thus the second stage of ligand exchange—ligand binding—fails to occur. In the case of ø-S and ø-L, this failure occurs because they follow P2-ø, during which P2 dissociates randomly. For steric displacement, this failure occurs because substrate displaces P2 before ligand does. Steric displacement is unique among the futile events in that both stages of nucleotide exchange take place without the corresponding ligand exchange process. For an interpretation of these futile paths in myosin, see SI 2.3.

The target cycle and futile events can also be described using eight ‘behavior blocks’. Behavior blocks are intervals of two or more transitions that capture important behavior. They are defined by the two states that lie on either end of the interval and the intermolecular reactions that take place between them (see SI 2.8). The three blocks that define the target cycle are: allosteric displacement 1(ad1), reverse (rev), and allosteric displacement 2 (ad2). Each futile event is its own behavior block. Hence, the eight behavior blocks are:

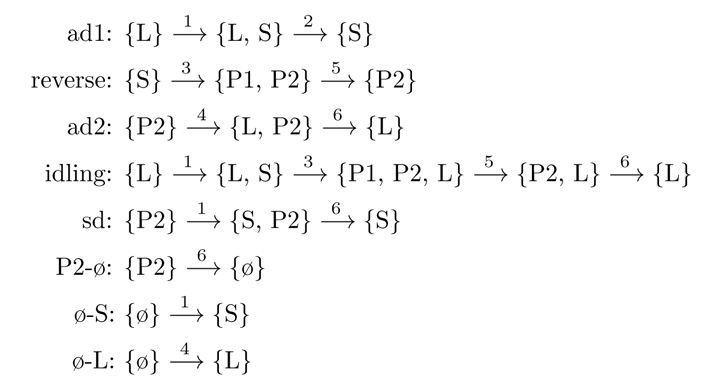

### Stochastic simulations

#### Constructing the single molecule reaction network

The geometric and node connectivity rules were used to construct a reaction network (see SI 2.4; states in which substrate or ligand are bound by their two outer nodes only are eliminated.). Each state in the network of 449 states has a different node configuration, eighteen of which are shown in Fig. 1D. As described, transitions between states can be made if two states differ by one more, or one less node connection, which corresponds to a forward and reverse reaction for any two states connected (by an edge) in the network. This transition rule produces six different linkage reactions, each of which can be mapped to a chemical equivalent that captures intermolecular, intramolecular or catalytic change (equivalent linkage transitions are stated on the right, where the words ‘connect’ and ‘disconnect’ refer to single node association changes):

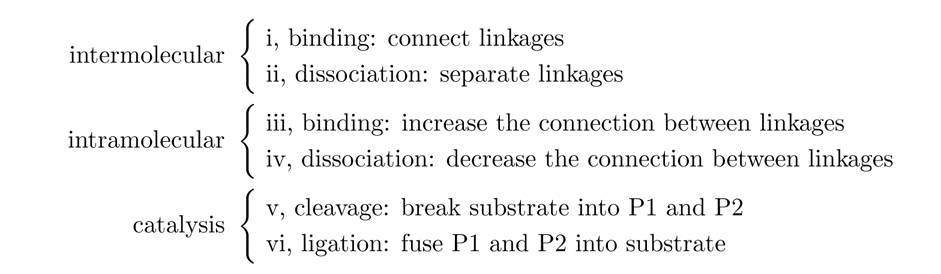

Using these six transition types to connect the states together results in a reaction network of 449 states, connected by 3449 possible transitions. The network represents all possible pathways in the time evolution of a single enzyme as it reacts with multiple copies of the four reactants S, P1, P2 and L. To reduce the number of rates required to perform simulations on the network, the following two simplifications were made. One, dissociation events at each of the six nodes were assigned one rate specific to that node (*k*_off-node_; e.g. *k*_off-a_), regardless of whether the reaction was intramolecular dissociation (reaction iv) or intermolecular dissociation (reaction ii). Two, intramolecular binding events (reaction iii) at every node were assigned the same rate (*k*_uni_). These two simplifications allow thirteen reaction rates to be used for simulations: four ‘concentration’ dependent intermolecular binding rates (reaction i; one for each of the four reactants: *k*_on-(S, P1, P2, or L)_); one intramolecular binding rate (*k*_uni_); six node dissociation rates (*k*_off-(a, b, c, d, e, or f)_); one cleavage rate (*k*_clv_); and one ligation rate (*k*_lig_) (see SI 2.5 for rate-labeled target cycle).

Values were assigned to the rates by following four guidelines. First, node energies were designed to be weak (no more than several kT), to allow reversible binding at each node association. Second, a thermodynamic hierarchy was set in which substrate binds the tightest, followed by ligand, followed by P2 and finally P1. This hierarchy constrains the respective sums of node energies (*ε*_node_) for each molecule according to:

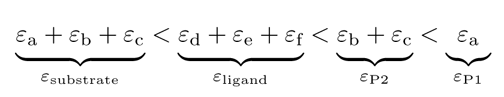

where negative energies reflect that substrate, as the most negative value, is the tightest binder (see SI 2.6). The six node dissociation rates were thus chosen according to this constraint, using local detailed balance to convert energies to rates (SI 2.7; eq. 9). Third, the intramolecular binding rate (*k*_uni_) was assigned a value of 1 10^6^ s*^−^*^1^, which is several orders of magnitude faster than the other rates in the system (see SI 2.4.4.ii). This assignment ensures that divalent binding is metastable. Most importantly, P2 binding is made metastable, which inhibits its random dissociation, or *{P* 2*} → {*ø*}*. And fourth, the cleavage and ligation rates were set equal to one another (as *k*_cat_), so that catalysis is inherently unbiased.

#### DIV control system, counting pathways, and categorizing futile pathways

Gillespie simulations were performed on the TRV linkage system, as well as on a control system (DIV), which has only one mechanical degree of freedom. Essentially, the control system is the catalytic side (left side, green) of the TRV system with the reactant molecules resized accordingly, such that the substrate and ligand bind divalently, and P1 and P2 bind monovalently. In table form, negative allosteric coupling in the DIV system compared to the TRV system is expressed like so:

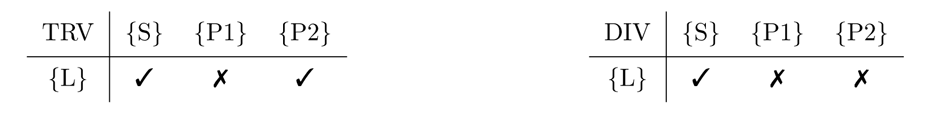

Thus, in the DIV system, S is allosterically coupled to ligand, but neither product is allosterically coupled to the ligand. The purpose of the DIV control system was to verify that, to emulate myosin, at least one product molecule needs to dissociate through an allosteric displacement, as does P2 in the TRV system.

Simulations were run as “single-molecule” experiments, in the sense that I only looked at the time evolution of a single enzyme as it reacted with the four reactants species. Analysis of the simulation results was a counting exercise. A series of simulations of the TRV and DIV systems were performed over a range of substrate concentrations, while keeping the ligand concentration fixed. Occurrences of the productive and futile pathways were then counted. Pathways were counted by matching the sequences of states preceding the release of each P2 molecule to pathway definitions (see SI 2.8). The counts were converted to enzymatic turnover rates, where the turnover rate is the number of P2’s released per second, and is used here as a measure of enzymatic activity. In addition to counts for the individual pathways, an overall turnover rate is also reported, which is the sum of all futile and productive activity.

In Fig. 2A the pathways are listed in legend form, where each pathway, or group of pathways is associated with the same color used in the plots. Fig. 2B is Fig. 1D redrawn using the eight behavior blocks: ad1, rev, ad2, idling, sd, P2-ø, ø-S, and ø-L. Using behavior blocks allows the information given in Fig. 2A to represent productive and futile pathways for either the TRV or DIV systems. Additionally, they allow the categorization of futile pathways. There are four categories of failure: idling, bad-ends, ligand-frees and bad-begins. Idling is idling as described above, bad-ends begin well (with ad1) but end wrong, bad-begins begin bad but end well (with ad2), and ligand-frees begin and end bad.

**Fig. 2.**
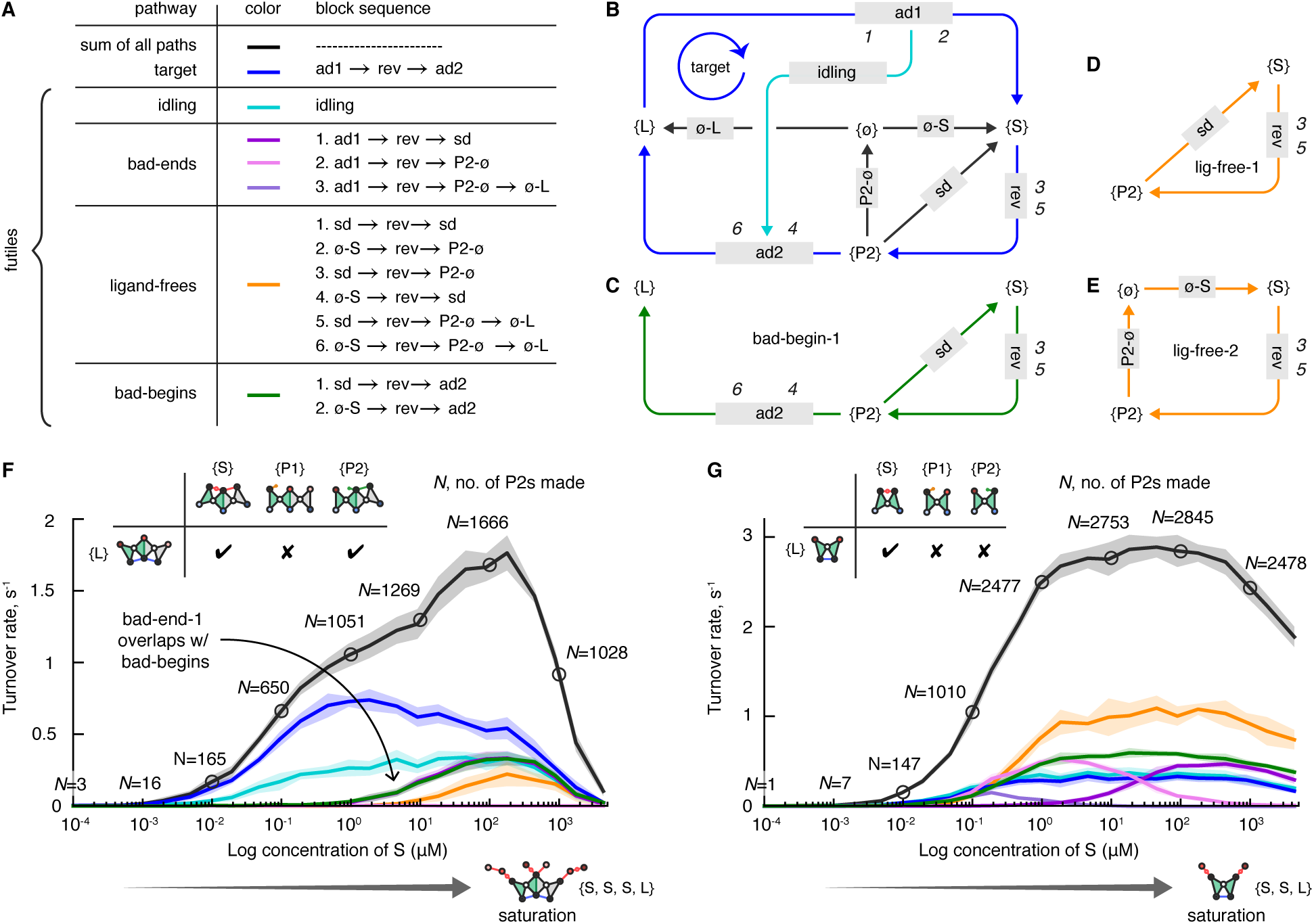
Simulation results. **A,** Pathway color legend with each pathway defined with its behavior blocks. **B,** Network showing the target and off-target pathways. **C-E,** Three examples of futile pathway subnetworks drawn using the legend color. **C,** bad-begin-1. **D,** ligand-free-1. **E,** ligand-free-2. **F,** Pathway analysis in the TRV system. The log scatter plot displays a total turnover rate (black) and one for each pathway, where the turnover rate is the rate at which new P2s are released from the enzyme. Substrate concentrations ranged from 0.1 nM to 5 mM and ligand concentration was held at 100 nM for all simulations. A total count of P2 release events (*N*) is given for every 10x change in concentration (circled points on black curve; see SI 2.9). The target pathway (blue) dominates for all substrate concentrations, though after 1 *µ*M, incidence of the futile pathways begin to rise. Overlap of bad-end-1 (purple) and bad-begins (green), and close-to-zero incidence of bad-end-2 (pink) and bad-end-3 (light purple), reflect the shortest futile path back to the target and low random release of P2, respectively (see main text). Around S = 100 *µ*M, the enzyme approaches saturation (‘saturation’ pic below), bringing the turnover rate back towards zero (see main text). **G,** Pathway analysis in the DIV system. Here, in contrast to the TRV system, the ligand-free pathways (orange) dominate for all [S] after 0.2 *µ*M, while the target cycle (blue) maintains a relatively low incidence. Also of note, bad-end-2 (pink), which reports on the random dissociation of P2, has a relatively high incidence at low [S]. As in the TRV system, saturation is approached around S = 100 *µ*M, though the effect is not as severe.

To show how individual pathways are embedded in Fig. 2B, subplots of bad-begin-1, ligand-free-1 and ligand-free-2 are shown Figs. 2C-2D, respectively, though individual pathways within these two futile categories are not distinguished in the plotted data.

#### Simulation results

A plot of the turnover rates for the TRV system shows how the behavior of the TRV system is dominated by the target cycle (Fig. 2F). At lower concentrations of substrate (below 100 nM), the target cycle occurs more frequently than occurrences of all the futile pathways summed together, verifying that the system operates as desired. At higher concentrations of substrate (above 100 nM), occurrences of the futile events begin to surpass the target cycle. This switch in behavior at higher concentration of substrate occurs because steric displacements of P2 by substrate become more probable at higher concentrations of substrate.

Diversions into the steric displacement pathway can be verified by identifying bad-end-1 as the primary pathway taken when the target pathway fails. This identification is made by noticing that, aside from idling, all diversions from the target cycle (or entries into a futile pathway) initiate at P2, which is the main branch point, and thus begin as bad-ends, all three of which are tracked individually. Because be2 and be3 do not occur, it can be concluded that the empty state (ø) is not visited, and all futile paths that go through the empty state (be2, be3, lf2-lf6 and bb2) do not occur; the only futile paths that need to be considered are idling, be1, lf1 and bb1.

With this simplification, two main diversions from the target cycle, which are combinations of a bad-begin-1, bad-end-1, and ligand-free-1, can be identified. The first takes place when a bad-end-1 is followed by a bad-begin-1, which is visible in the plot as overlapping curves. They occur together because they form the shortest combined pathway back to the target pathway if a single steric displacement (sd) takes place:

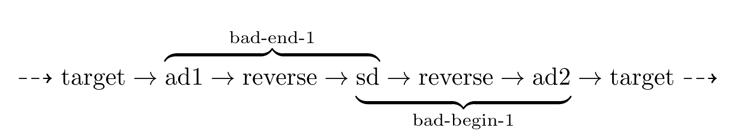

If two consecutive steric displacements take place, the second diversion can be identified as bad-end-1, followed by ligand-free-1, followed by bad-begin-1:

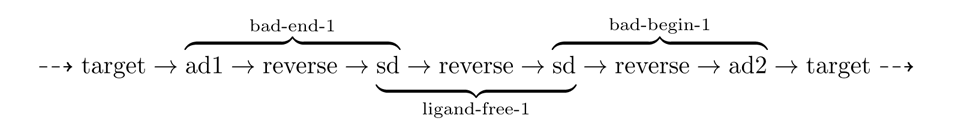

Inhibition of this sequence is visible on the plot as a lower turnover rate for the ligand-free pathways, which is really only ligand-free-1.

The mechanism by which these two principal diversions, consisting of one or two steric displacements, are inhibited is described in SI 2.10. The main point to be made is that allosteric displacement of P2 by ligand (ad1) wins out over steric displacement by substrate (sd) because the first ligand that binds can complete its displacement of P2, whereas multiple attempts by multiple substrates are required for a successful steric displacement. This difference is due to the metastable two-node association (at nodes e and f) ligand immediately makes with the enzyme, compared to the unstable single-node association (at node a) substrate is restricted to when P2 is bound. Consequently, steric displacement is only possible at very high [S], where the high incidence of substrate binding increases the probability of success.

Incidence of idling follows a different pattern than the other four futile pathways (Fig. 2F, turquoise line). Idling is less sensitive to changes in substrate concentration, and though its occurrence steadily rises with a rise in substrate concentration, it is not “activated” at certain substrate concentration as are the other futile pathways. Eventually, the incidence of idling decreases along with the overall enzymatic activity of the system, as the system reaches saturation and then inhibition, which is described next.

The total turnover rate with ligand (Fig. 2F, black line), which is the summation of the target and futile pathways, increases with the concentration of substrate, until around S = 100 *µ*M where saturation is reached and an increase in substrate leads to a sharp decrease in enzymatic activity. Accordingly, all the contributing turnover rates also decrease at high [S]. By visually inspecting trajectories for simulations done at high [S], I infer that the decrease begins to take place as the substrate’s binding rate from solution approaches the intramolecular binding rate. As this happens, the enzyme continually saturates with one ligand and up to three substrate molecules, which makes it difficult for a single substrate to displace the ligand from the enzyme and form the cleavage complex, and consequently the turnover rate plummets (see S?).

In contrast to the TRV system, behavior of the DIV system is swapped (Fig. 2G). The DIV system is dominated by the occurrence of the ligand-free pathway (orange), as well as the other futile pathways emerging from the P2-bound state (P2). Incidence of the target pathway (blue) remains relatively low. In comparing the DIV and TRV systems, it is important to keep in mind that in the DIV system, steric displacement (sd) and allosteric displacement (ad2) of P2 are not displacements as they are defined for the TRV system. In the DIV system, P2 is bound only monovalently, and thus it dissociates randomly. Hence, occurrences of pathways that include the sd and ad2 behavior blocks (which is all pathways except for ligand-free-2, bad-end-2 and bad-end-3) reflect only that substrate, or ligand happened to be bound to the enzyme when P2 dissociated.

That P2 dissociates randomly in the DIV system is jointly reflected by zero incidence of bad-end-1 (purple), and visible incidence of bad-end-2 (pink) and bad-end-3 (light purple), at low concentrations of substrate. Bad-end-1 is exclusive to one pathway that reports on sd (ad1 → rev → sd), or the presence of substrate when P2 dissociates. Hence, its zero incidence at low [S] means that substrate is not present when P2 dissociates. Likewise, the relatively high incidence of bad-end-2 and bad-end-3 at low [S], both of which include the P2-ø block, which is the transition from the P2 bound state to the empty state, confirms that P2 dissociates randomly. Note that bad-end-2 and bad-end-3 have a near zero incidence in the TRV system, as P2 is almost always displaced by ligand in the TRV system.

As the concentration of substrate rises in the DIV system, and it becomes more probable for substrate to bind to the empty enzyme, and bind to the enzyme before P2 dissociates (at the empty P1 site), two effects become visible on the plot. The first effect is that the brief incidence of the bad-end-3 pathway (light purple), which reports on ligand binding to the empty enzyme (L-ø), decreases to zero as substrate binding to the empty enzyme overtakes ligand binding to the empty enzyme. Coincidentally, the ligand-free and bad-begin curves rise (between 0.1 and 1 *µ*M), as they both include pathways that report on substrate binding to the empty (S-ø). The second effect that becomes visible is that incidence of the bad-end-2 pathway (fuchsia), which reports on the random dissociation of P2 leading to the empty state, decreases to zero. This decrease coincides with an increase in incidence of the bad-end-1 pathway (dark purple), which reports on substrate binding to the enzyme before P2 dissociates (sd).

In the DIV system, occurrences of *idling* more or less tracks with occurrences of the target cycle, whereas in the TRV system idling events happen significantly less than that of the target cycle (Fig. 2F & 2G, compare blue and turquoise lines). This difference further reflects better performance of the TRV system, though the reason for this difference is not completely clear.

Saturation and inhibition also take place in the DIV system, as can be seen in the decrease in the total turnover rate at high substrate concentration (See SI 2.12 for a calculation of the maximum turnover rate in the DIV system).

In the supplement, I give three additional presentations of the simulation data (SI 2.13). The first plot compares the ability for ligand to stimulate enzymatic activity in each system, using data from simulations performed with and without the presence of ligand (SI 2.13, Fig. 17A). The plot shows how enzymatic activity is highly dependent on the presence of ligand in the TRV system, whereas it is not in the DIV system. This ligand-activated enzymatic activity, in the TRV system, reproduces actin-activated ATPase activity in myosin [19]. The second plot compares target cycle efficiency in each system (SI 2.13, Fig. 17B). The third is a plot of an individual trajectory of the TRV system at “peak” performance (SI 2.13, Fig. 18).

### Constructing a dimeric walker

#### Generating a landing bias

In this section, mechanical behavior is extracted out of the system to construct a myosin-like dimeric walker (Fig. 3). To do this construction, I use the rigid states (*C*_1_*, C*_2*A*_*, C*_2*B*_*, C*_3_ and *C*_4_) defined in Fig. 1D. The goal is to “derive” a condition for directed walking on a track that is consistent with the logic of the linkage system and uses what is known about walkers.

**Fig. 3.**
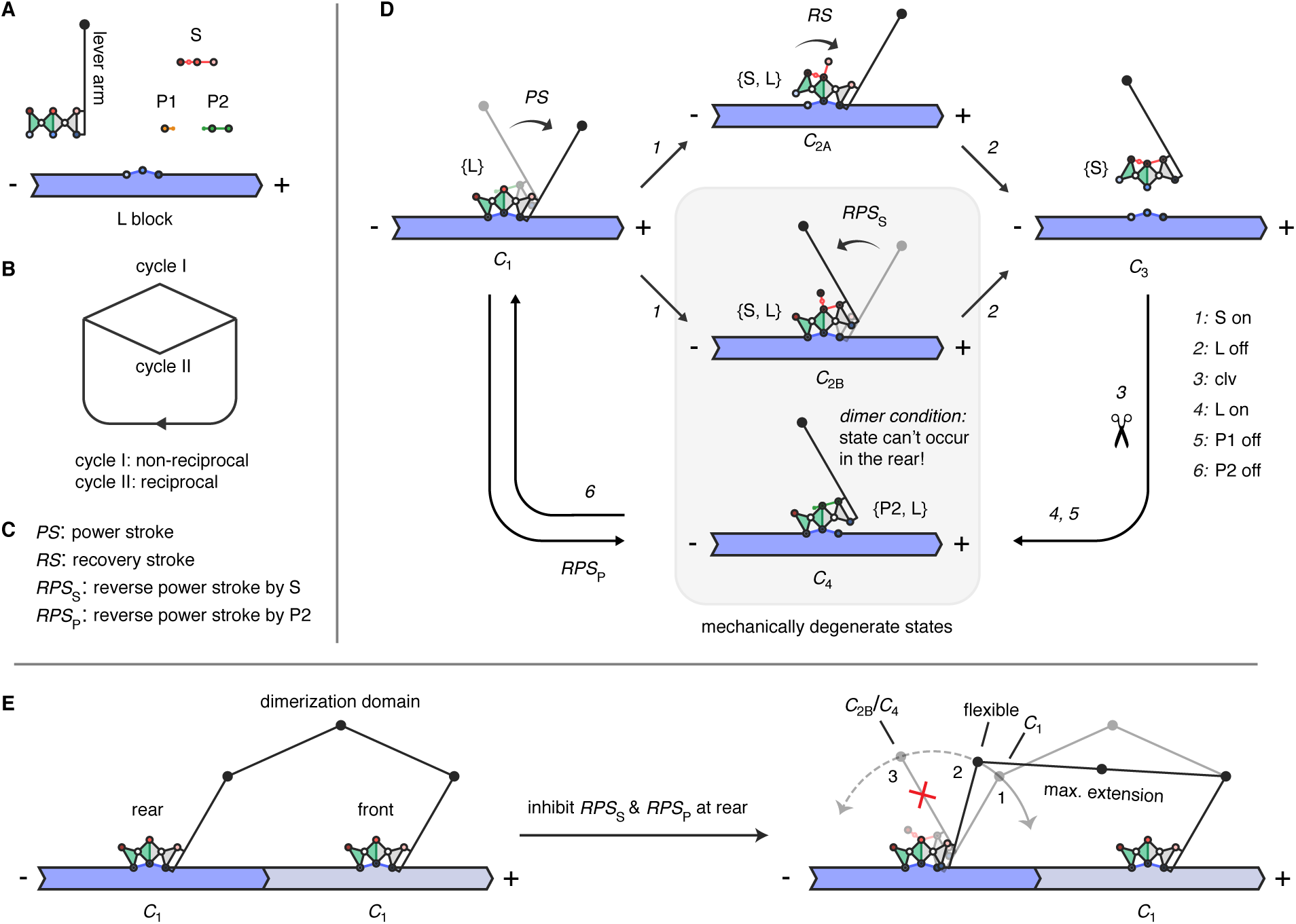
Two track cycles and a dimeric configuration that reinforces the non-reciprocal one. **A,** Five molecules of the linkage track system. There are two changes made here: (1) a lever arm is rigidly attached to the right-hand side of the enzyme at two nodes; and (2) the ligand is transformed into a block-like polar molecule, which contains the same trivalent binding site, except now it held rigidly in the block. Polarity is indicated by the (-) and (+) signs, where (+) is the desired direction of motion. **B,** Mechanical cycles key **C,** Reactions key. **D,** Mechanical cycles. Only the rigid states in Fig. 1D are displayed here in the track version of the system. Along mechanical cycle I (starting at *C*_1_/*{*L*}*), substrate binding on the left side of the enzyme (*RS* and *C*_2*A*_) allows the lever arm to maintain its forward orientation during detachment to state *C*_3_. Following cleavage, the enzyme reattaches in state *C*_4_, in which the lever arm is pointed towards the minus (-) end of the track. In the final transition, the power stroke (*PS*) takes place, returning the system to state *C*_1_. Along mechanical cycle II, substrate binding on the right, causes detachment through state *C*_2*B*_ and the *RPS_S_*. Reattachment is the same as it is in mechanical cycle I, however, *C*_4_ is mechanically degenerate with *C*_2*B*_ (gray box), leading to the dimer condition, which says these two states cannot occur in the rear position of a dimer (see main text). **E,** Satisfying the dimer condition. Left, dimeric construct with both units tightly bound in state *C*_1_, and the dimerization domain (two extra bars) relaxed. Right, upon dissociation of f-f* on the rear unit and counter-clockwise rotation, the dimerization domain reaches its maximum extension at some angle and flexible state between *C*_1_ and *C*_4_/*C*_2*B*_, which in turn inhibits (red ‘x’) the *RPS_S_* and *RPS_P_* from taking place.

A simple framework says that directed motion in a dimeric walker arises in two ways [17, 18]:

1. A landing bias: in which a unit that dissociates is biased to land at the front binding site.
2. A pickup bias: in which the rear unit is biased to detach more than the front unit.

In the walker described, a landing bias is generated through allosteric feedback between two linkage units, transmitted when both units are bound to their track. The track system in constructed by altering the monomer system in two ways (Fig. 3A). One, a lever arm is rigidly connected to the right-most triangle of the enzyme. Two, the ligand is converted into a polar block-like molecule that has an extended structure, containing a single binding site. The trivalent binding site allows oriented (stereospecific) binding to the track, as in myosin [14, 20].

By considering only the five rigid states in the target cycle and leaving out the intervening states, two mechanical cycles emerge (Fig. 3D; 3B & 3C are legends), mechanical cycles I and II. Mechanical cycle I is non-reciprocal (it obeys time-reversal asymmetry) and mechanical cycle II is reciprocal (it is time-reversal symmetric)(see SI 2.14). The difference between the two cycles results from two different track detachment pathways, or two different pathways along which substrate can displace ligand.

The different mechanical behaviors of the lever arm along each cycle are characterized in this way: After substrate loading and detachment from the track (ad1), the angle of the lever arm is reverted towards the minus end (-) of the track, with respect to the center line of the enzyme when detached. Along mechanical cycle I, this reversion is allowed to take place after detachment, as a diffusive process, because substrate first binds divalently on the left side of the enzyme, which does not change the angle of the lever arm (Fig. 3D, *C*_1_ → *C*_2*A*_). This detachment pathway is named the recovery stroke (*RS*). Along mechanical cycle II, the reversion must take place while the enzyme is still attached, because substrate first binds divalently on the right side (Fig. 3D, *C*_1_ → *C*_2*B*_). This pathway is thus named the *reverse power stroke* (*RPS_S_*), where the ‘S’ subscript denotes substrate ^1^.

During attachment to the track, and the exhausting of P2 (ad2), the angle of the lever arm rotates towards the plus end (+) of the track. This stage is thus named the power stroke (*PS*) (Fig. 3D, *C*_4_ *→ C*_1_).

To generate a landing bias, a *dimer condition* is invoked, which says that *C*_4_, the P2-bound and pre-power stroke state that occurs upon re-attachment to the track, cannot occur in the rear position of a dimeric construct. The logic used to invoke the condition is this:

Since the transition from *C*_3_ to *C*_4_ is reattachment to the track, assuming the other unit in the dimer is attached to the track, the only way to get forward motion is for the detached unit to reattach in front.

Because *C*_4_ and *C*_2*B*_ are mechanically degenerate, the dimer condition automatically rules out detachment from the track through *C*_2*B*_ at the rear unit. Consequently, a mechanism is required by which a front unit inhibits occurrence of mechanical cycle II and *RPS_S_* in a rear unit. By symmetry, the mechanism will also inhibit reversion of the lever arm if P2 binds at the rear (Fig. 3D; going in reverse from *C*_1_*, C*_1_ *→ C*_4_). A reversion caused by P2 is also named a reverse power stroke (*RPS_P_*), where the ‘P’ subscript denotes that P2 rather than substrate causes the reversion.

As shown in Fig. 3E, satisfying the dimer condition is arranged using a dimerization domain that limits counterclockwise rotation of the right side of the rear unit, so that substrate or P2 cannot divalently bind on the right side, between b-b’ and c-c’. The length of the dimerization domain is such that at some rotated angle of the rear unit’s lever arm, lying between *C*_1_ and *C*_2*B*_/*C*_4_, and accessible when f-f’ dissociates and the right side becomes flexible, the dimerization domain will reach its maximum extension. From this flexible state, the unit can only return to *C*_1_, and rigid trivalent track binding when f-f’ reforms. The assumption being made here is that the front unit remains tightly and rigidly bound when substrate or P2 bind to the rear.

#### How a forward step is made

There are four different pathways (binding cases) by which substrate can bind to the dimer. The *1st* binding case was described in the previous section, where substrate attempts to bind divalently on the right side of the rear unit, but cannot cause detachment due to the inhibitory mechanism. The three other binding cases to describe do lead to track detachment, and thus motion of the dimer units: the *2nd*, divalent binding of substrate starting on the left side of the rear unit, or through *C*_2*A*_; the *3rd*, divalent binding of substrate starting on the right side of the front, through *C*_2*B*_; and the *4th*, divalent binding of substrate starting on the left side of the front, through *C*_2*A*_. The *2nd* case leads to a forward step of the dimer (Fig. 4A). The *3rd* and *4th* binding cases lead to what is called a foot stomp in the myosin literature [25], in which the front unit detaches and reattaches in front (Fig. 4B).

**Fig. 4.**
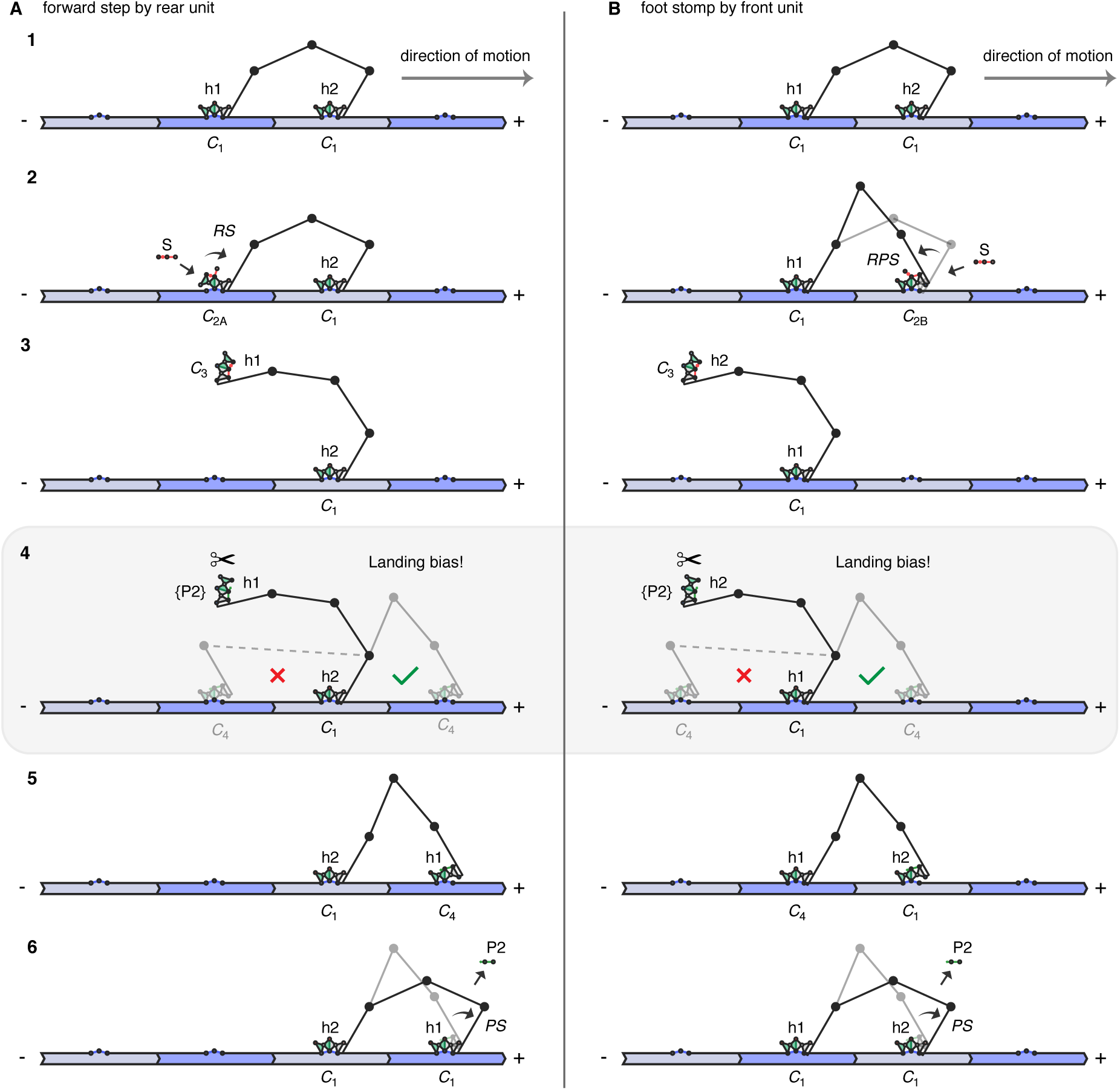
A forwards step and foot stomp of the walker. **A,** A forward step. In steps 1-3, because of feedback from the front unit, the rear is displaced from the track through the recovery stroke pathway (mechanical cycle I). In step 4, after cleavage, the detached P2-bound unit is subject to the landing bias. The landing bias is a consequence of the feedback mechanism used to satisfy the dimer condition (see Fig. 3E), which says that *C*_4_ cannot occur in the rear. A forwards step is completed in steps 5 and 6, ending with the power stroke at the front unit. **B,** A foot stomp. In steps 1-3, because there is no inhibitory feedback on the front unit, the front unit is displaced from the track through the *RPS_S_*pathway (mechanical cycle II); though either detachment pathway is possible. In step 4, because of symmetry, the same situation arises as for the rear unit—after cleavage, the detached P2-bound unit is subject to the landing bias. A foot stomp is completed in steps 5 and 6, ending with the power stroke at the front unit, but no forward progress.

A forward step (*2nd* binding case) begins when substrate binds and displaces the rear unit from the track through *C*_2*A*_, along mechanical cycle I (Fig. 4A, steps 1-3). After cleavage and dissociation of P1—which leaves the detached rear unit bound to P2, flexible and thus able to rebind the track—the inhibition of state *C*_4_ in the rear position becomes important to generating the forward landing bias (step 4). The bound rear state (left side and opaque) is the inhibited *C*_4_ state (indicated by a red ‘x’) shown in Fig. 3E. It can’t occur because the dimerization is not long enough to span the distance (indicated by the dashed line). However, binding to the front site (right side and opaque) is permitted because the dimerization domain can easily span the distance between the ends of the lever arms (indicated by green check mark). After attachment at the front, the power stroke takes place, which displaces P2 to reform the starting dimer configuration, but progressed one site in the forward direction.

A foot stomp begins when substrate binds and displaces the front unit, through either *C*_2*B*_ (*3rd* binding case) or *C*_2*A*_ (*4th* binding case). By contrast to the rear unit, the front unit is not inhibited from going through mechanical cycle II, because counterclockwise rotation decreases the distance between the lever arms. P2 can also rebind to the front unit, to form *C*_4_, but will likely be re-displaced before causing detachment. Upon detachment by substrate binding, the exact same situation arises as for detachment at the rear unit—a landing bias is generated after cleavage. Hence, just like for the rear unit, after reattachment at the front, the power stroke takes place, which displaces P2 to reform the starting dimer configuration, but in this case no forward progress has been made.

The walker is a directed walker due to a landing bias alone, which causes forward motion largely by preventing backwards motion. The mechanism is related to mechanisms proposed for myosin, in which the recovery stroke causes a penalty for binding at the rear position [26, 27]. There is no pick-up bias in the system because there is no mechanism that inhibits substrate binding to the front unit—the *3rd*, and *4th* binding cases are easily accessible. Substrate binding is only inhibited along one of two binding pathways to the rear unit (the *1st* binding case). Thus, if anything, there is a negative bias that favors detachment of the front unit, which increases the inefficient use of substrate. In other words, the landing bias system described is in principle a directed walker, but an inefficient one.

In myosin, a pick-up bias (referred to as gating) is believed to come from intermolecular strain between the units of the dimer that allows the front unit to retain ADP longer than the rear unit [28, 29]. Similarly, a pick-up bias can be created in the linkage system with tighter coupling between the units (see SI 2.15, pick-up bias walker). At the other extreme, a walker with no directional bias can also be created (see SI 2.15, no-bias walker). In the linkage system, the pick-up bias is not a mechanism for generating directionality, rather, just as proposed for myosin [18], it is in principle a mechanism for generating efficient directed motion.

## Discussion

### A geometric description of directionality

In addition to satisfying the dimer condition described above, an alternative geometric interpretation of directionality can also be described. The geometric interpretation says that a landing bias and directed motion is possible if the reciprocal cycle is inhibited in the rear unit, by using allosteric feedback to select the non-reciprocal cycle. The goal is to control the path taken during the recovery stroke phase (inhibit *C*_2*B*_ in the rear). Hence, within this geometric interpretation, rather than directly controlling for landing, the landing bias emerges from using feedback to control for non-reciprocity.

Non-reciprocal conformational cycles are known to be important for motility at the microscale (the “scallop” theorem) [30]. The scallop theorem says that because motion is inertialess at the microscale, microscale organisms and machines cannot propel themselves by a symmetric and reversible change of shape, as the opposing changes will cause movements that cancel each other out. To break directional symmetry and ‘swim’, they must push against the medium in a continuous and cyclic sequence of non-reciprocal shape changes.

As motion is also inertialess at the nanoscale, it has been proposed that molecular motors also use non-reciprocal conformational changes to produce directed motion [31–35]. However, molecular motors differ from microscale machines in two important ways. One, they operate stochastically—their reactions are thermally activated and highly reversible [35, 36]. Two, they ratchet progress along track-like structures [37–41], making use of two or more enzymatic units [42–44]. Stochasticity means that continuous conformational change along a non-reciprocal cycle cannot be assumed [33]. The continuous change assumed for microscale motility is the time-averaged activity of many internal molecular motors working in concert, as in flagella [45], or cilia [46]. The linkage system suggests three ideas about how non-reciprocal cycles may help individual molecular motors achieve directionality in track systems, in particular, when the motors and tracks are not topologically linked, as they are in catenane systems [47]. One, it suggests that monomer units may complete a chemical cycle through reciprocal or non-reciprocal conformational cycles, due to the symmetry involved in allosteric communication, and inherent stochasticity. Two, it suggests that non-reciprocal cycles can break directional symmetry when oriented along a track-like structure, though this instance can be erased by unfavorable binding events. Three, it suggests that for symmetry breaking to not be erased, the role of a second unit, which is further along in the motor cycle, might be to help reinforce non-reciprocity in the lagging motor unit, through allosteric feedback.

In [33] and [34], respectively, similar concepts to ideas one and three above are explored, but for stochastic swimming rather than motion along a track.

### Relevance to engineering molecular motors

The chain-like linkage geometries used here are simple compared to the complex three-dimensional geometries of proteins and ribonucleoproteins. Hence, only a small space of allosteric behavior is explored, which is negative allosteric cooperativity between two binding sites (pairwise cooperativity). Even in longer versions of these chains, events happening at any single “link” in the chain can only affect adjacent links, similar to how DNA strand displacement progressively takes place [48, 49]. This attribute places limitations on the complexity of allosteric signalling that can be achieved beyond pairwise cooperativity, which is a large space [6, 50, 51].

For example, in modeling myosin, important behavior was left out, the most significant being that ligand (actin) binding cannot trigger dissociation of the P_i_ analog P1, or trigger lever arm movement coupled to this dissociation. In myosin, actin binding triggers the release of both products, and both releases are coupled to lever arm motion. Adding a third binding site that positively or negatively affects allosteric coupling between two other sites would also be difficult with chain-like structures.

Including these additional allosteric behaviors likely requires extending the model to two dimensions. Two-dimensional mechanical networks have been explored as allosteric materials in several studies [52–56]. These networks have multiple independent degrees of freedom that allow complex mechanical responses. However, within a chemical context, it is not clear how these complex internal responses can be triggered, or recovered once created. Internal mechanical responses must be connected to external responses to transduce chemical information. Along these lines, chemical to mechanical signal transduction has been experimentally demonstrated in DNA mechanical networks, whereby DNA signals were used to trigger uniform conformational change in ‘accordion’ networks, which have a single degree of freedom [57, 58].

Using two mechanical degrees of freedom, the linkage system models allosteric signal transduction by representing mechanical networks as agents that interact through multivalent contacts—agents that join together and come apart in discrete steps that dynamically affect the conformation of the connected structure. The result is a switch-like reaction (allosteric displacement) that allows an incoming chemical signal to be converted into an internal mechanical signal, and released as a different outgoing chemical signal. Integrating this concept into two and three-dimensional networks may be possible.

Cyclic activity was demonstrated by connecting two displacement reactions together, by catalysis [59]. The catalytic cycle of the TRV system shows how completion of a fuel cycle (fuel binding, cleavage, and product dissociation) can be made dependent upon a second process (ligand binding and dissociation). Both processes must take place to efficiently complete the cycle [7, 60, 61]. Allosteric coupling is maintained by joining the two such that the end of each process (energetically uphill dissociation) overlaps with and is initiated by the beginning of the other (downhill binding). As demonstrated by the DIV system, uncoupling product dissociation from ligand binding results in cycles dominated by the futile use of fuel.

In [60, 61], it is argued that mutually dependent allosteric couplings are at work in most biomolecular machines. The expectation is that these types of mutual couplings will also be important in the design and construction of synthetic and autonomous molecular machines [60]. Here, the argument is made using the linkage system, which shows two-way allosteric coupling and the controlled cycling it can produce. Although allostery is not required to construct synthetic molecular motors, it will likely be important in the design of molecular motors that operate in mixed-purpose teams, and use the same chemical gradient as fuel.

The linkage abstraction ideally represents concepts that can be realized using multiple synthetic approaches. In that sense, one can imagine using DNA or RNA [62–65], rotaxane/catenane chemistry [4], or peptide chemistry [66, 67] to construct systems of interacting linkages. A major challenge in studying biomolecular machines is figuring out which structures are essential to the function they demonstrate, and which may be nonessential [68–70]. By developing allosteric models that generate the internal steps required to achieve various intermolecular outcomes, we can better mimic biomolecular motors through interpolation, without knowing exactly how they may operate. The linkage model is intended to be a contribution towards creating such methodologies.

## Acknowledgements

I thank Zev Bryant and David Soloveichik for many helpful discussions about the work. In addition, I thank Dean Astumian, Keenan Breik, Miranda Holmes-Cerfon, Rizal Hariadi, Robert Kohn, Muneaki Nakamura, Enrique Rojas, Howard Stone, and Erik Winfree for feedback about the ideas within. This work was supported by a National Institutes of Health (NIH) Fellowship F32GM09442 to Tosan Omabegho, while a postdoctoral scholar at Stanford University.

## Footnotes

1 In the myosin literature, the ‘recovery stroke’ refers to the reversal of the power stroke (or repriming) triggered by ATP binding. It is believed to take place after actin detachment, during tight ATP binding [21–23], or hydrolysis [24]. Since there are two distinct substrate binding pathways in the linkage system, I use the term recovery stroke (*RS*) only for the pathway in which the lever arm is not required to reverse before track detachment—the *RS* pathway. Along the other substrate binding pathway, the lever arm is required to reverse before detachment—the *RPS_S_* pathway. Hence, I use ‘reverse power stroke’ (*RPS*) to mean a reversal of the lever arm that takes place while the motor head is bound to its track, which can be caused either by substrate binding (*RPS_S_*) or P2 binding (*RPS_P_*).

2 *k*_on-S_, at 5 mM [S], is calculated by multiplying the bimolecular rate constant, *k*_bi_ = 9 × 10^6^M^-1^s^-1^, by 5 mM

## 2 Supplemental

### 2.1 Displacement of ligand by substrate

**Fig. 5.**
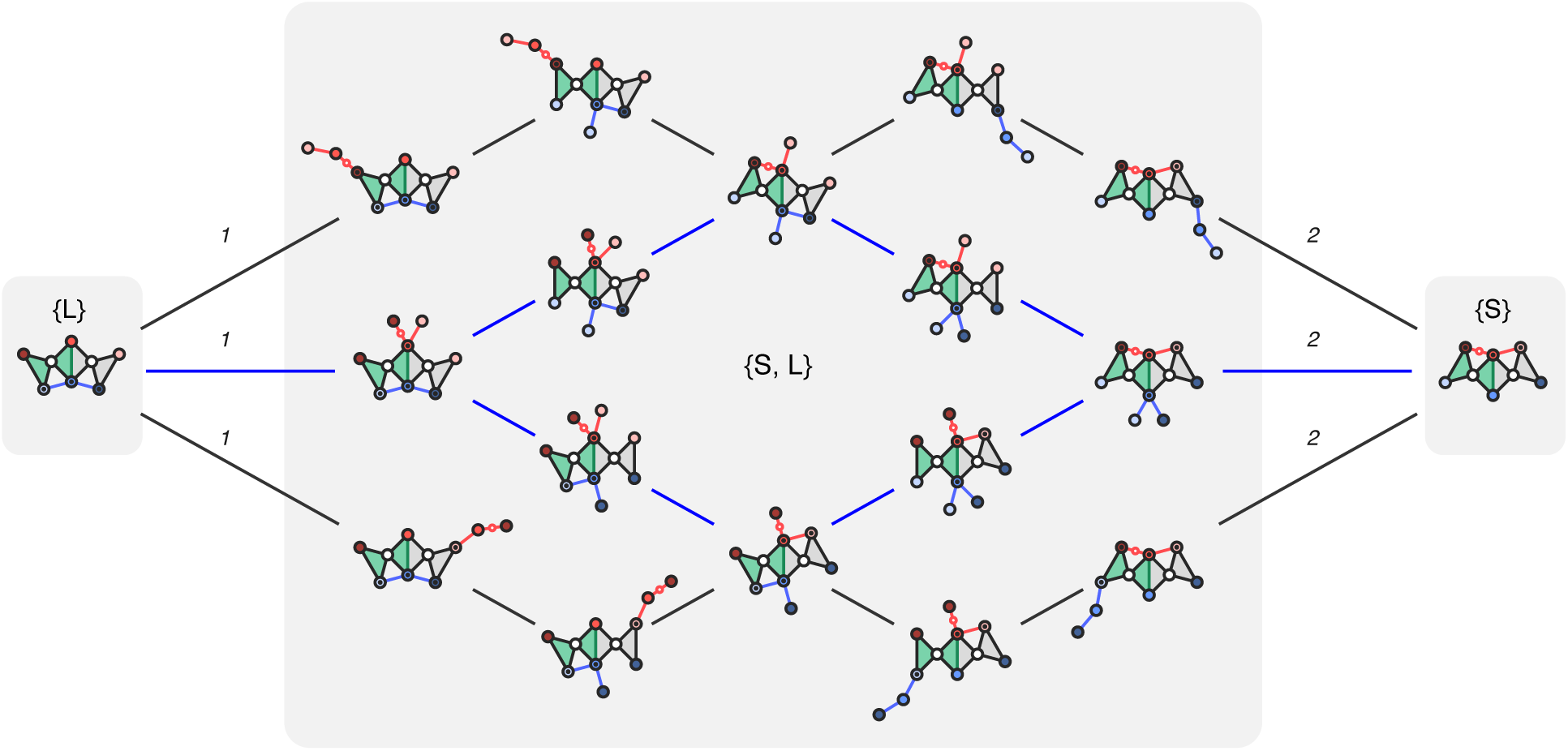
Network representation of S allosterically displacing L. The two inner paths (blue) are the pathways shown in Fig. 1D in the main text. Note that this is still a partial network of displacement, which represents the most probable paths given that intramolecular binding is much faster than intermolecular binding than dissociation (*k*_uni_ ≫ *k*_off_’s). The complete set of pathways include those in which consecutive in which consecutive intramolecular dissociation events take place before intramolecular binding. For example, if after substrate bound to node a, the ligand were to dissociate from nodes d and e, one after another, before substrate bound to b. Intermolecular reactions labeled: 1, S on; and 2, L off. These two reactions apply to the forward direction of the reaction sequence, going from left to right; though all transitions are reversible.

### 2.2 Complete futile event pathways

### 2.3 Futile paths in myosin

A biochemical interpretation of idling, and the random loss of product (‘P2-ø’) in myosin, are given here. These pathways would arise as a result of nucleotide exchange (the driving process) getting out of sync with actin exchange (the driven process).

#### idling

In the event that idling happens in a myosin dimer system, a rear myosin head (the head which supposed to advance) will remain bound to actin through ATP binding and hydrolysis. As a result, the head will not detach and will not advance, though one molecule of ATP has been hydrolyzed. Thus, it can be said that idling leads to the futile use of ATP.

#### dissociation of Pi and ADP before binding actin (equivalent to P2-ø)

If a myosin monomer is able to easily dissociate from Pi and ADP before binding actin, this in principle means that myosin can go through a whole hydrolysis cycle without interacting with actin, which is by definition a futile use of fuel, as the hydrolysis cycle is not driving another biochemical process. Hence, coupling the loss of product to actin binding is essential to myosin’s ability to couple ATP processing to its mechanical activity on actin.

Furthermore, in the context of a myosin dimer, the presence of ADP on the actin-bound leading head is believed to act as a steric block that inhibits ATP from binding to the lead head while the trailing head binds ATP and is displaced from actin. Hence, in the case of the dimer, the coupling of actin binding to ADP dissociation helps keep the heads out of phase with one another, so that ATP can be used efficiently.

### 2.4 Constructing the reaction networks

The reaction networks were constructed in these four steps:

1. Generate basis states.
2. Map exclusion rules between basis states.
3. Generate complete set of states.
4. Connect states to one another.

Each of these four steps are described in detail below:

**1. Generate basis states.** The basis states were created by enumerating all the ways the single reactants can bind to the enzyme. For the DIV system there are nine basis states including the empty state (SI Fig. 7), and for the TRV system there are seventeen basis states including the empty state (SI Fig. 8). The TRV basis states excludes two geometrically possible states by invoking an ad hoc principle, named the *adjacency rule*, that only adjacent multivalent bonds can form (SI Fig. 9).

**Fig. 6.**
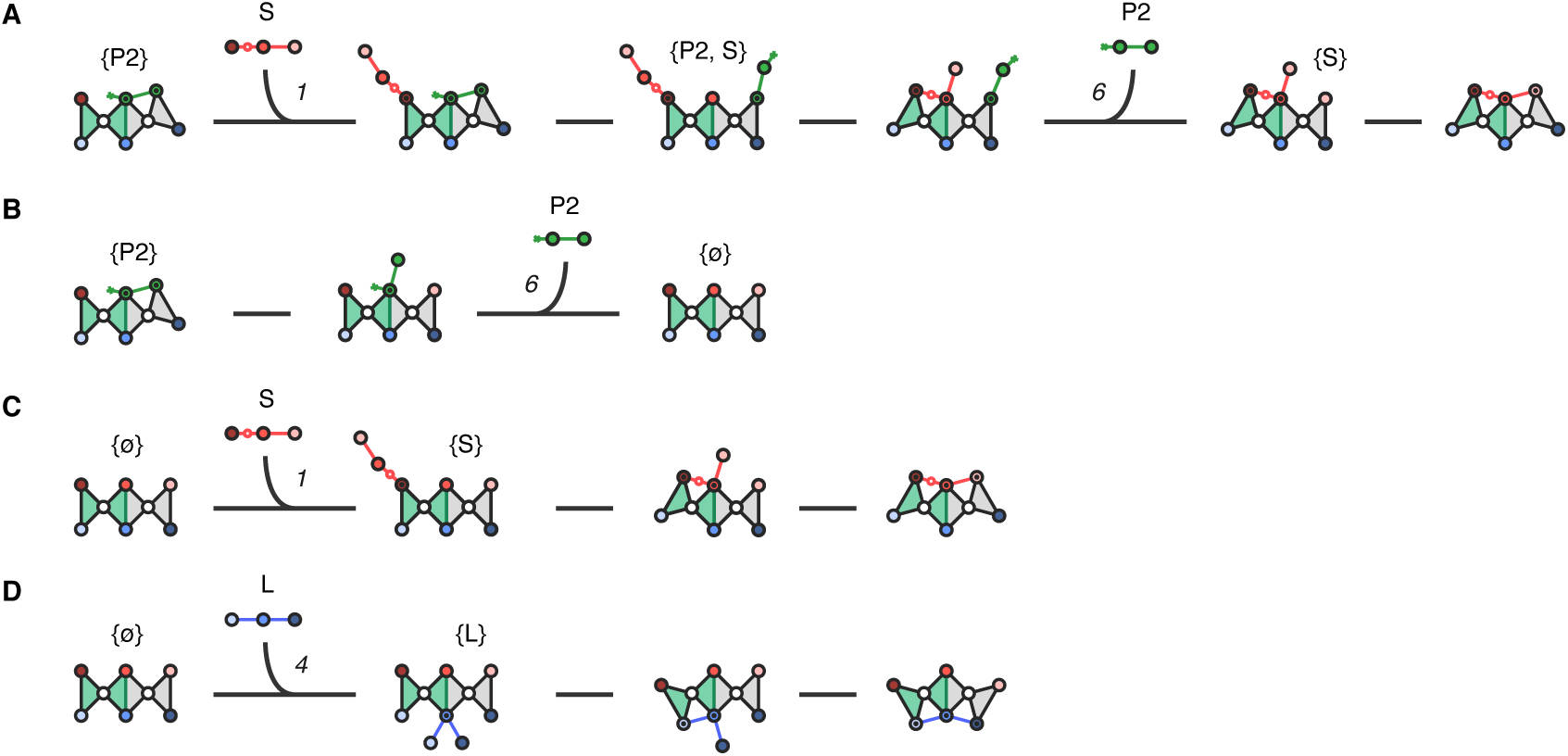
Completed futile pathways. These are complete versions of the abbreviated pathways shown in the main tex. A, Steric displacement. B, P2-ø. C, ø-S. D, ø-L. Intermolecular reactions shown: 1, S on; 6, P2 off; and 4, L on.

**Fig. 7.**
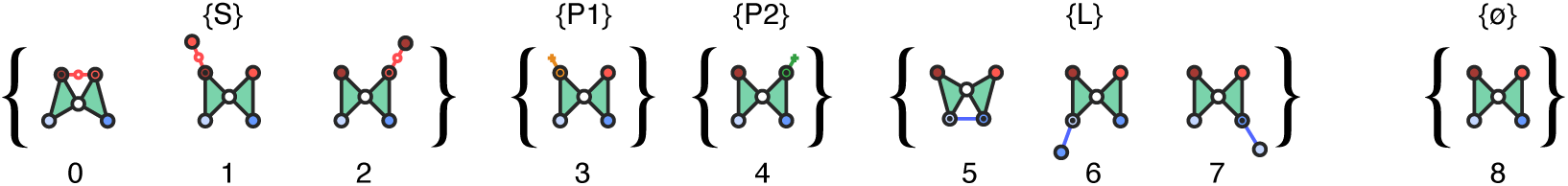
DIV bases states. The subset of eight basis states of the DIV system, which includes the empty state (ø). In each of these states (save ø) only one molecule of S, P1, P2 or L is bound, and the subset of states enumerates all the different ways these four molecules can bind to the enzyme.

**Fig. 8.**
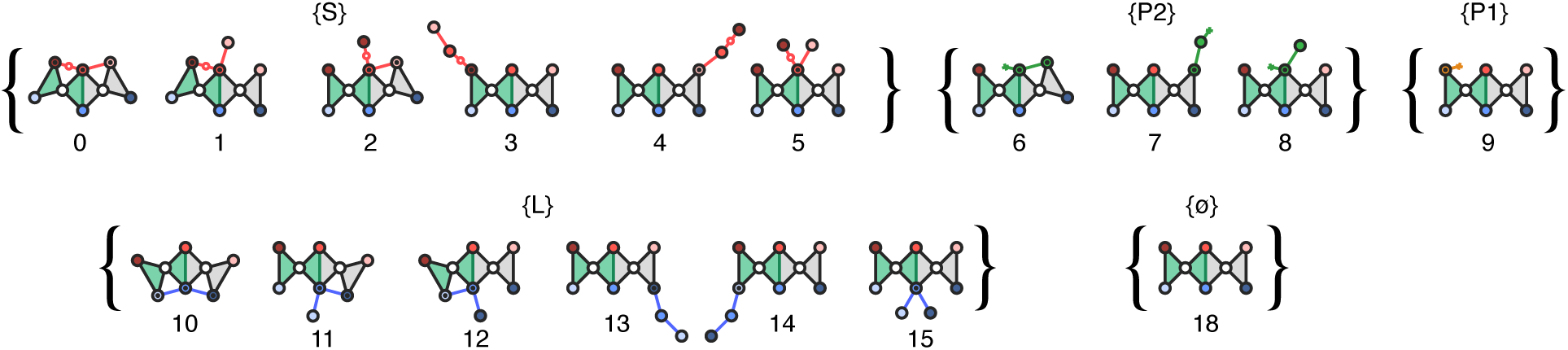
TRV basis states. Subset of seventeen states that are used to generate and name the complete set of 449 states. In each of these states only one molecule of S, P1, P2 or L is bound, and the subset of states enumerates all the different ways these four molecules can bind to the enzyme, barring the two states eliminated by the adjacency rule.

**Fig. 9.**
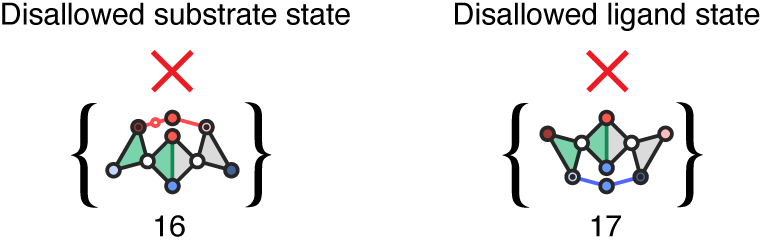
Adjacency rule. Only adjacent bonds are allowed to form or dissociate between two interacting linkages. Left, disallowed substrate state in which the two outer nodes are bound, and the central node is dissociated. Right, disallowed ligand state in which the two outer nodes are bound, and the central node is dissociated.

#### 2. Map exclusion rules between basis states

The exclusion rules for each system are mapped out as a binary matrix that matches the three sets of substrate and product basis states against the set of ligand binding basis states. For each matchup, the two states can either coexist together on the enzyme, and thus are an allowed two-molecule-bound state, or they cannot coexist together on the enzyme, and thus are an excluded two-molecule-bound state (symbolized by). What should be noticed is that excluded states only result from matching one multivalent state to another multivalent state. Thus, the DIV system has only one excluded state (*{*0, 5*}*), and the TRV system has nine excluded states: seven for S and L (*{*1, 12*}*, *{*0, 12*}*, *{*1, 10*}*, *{*0, 10*}*, *{*2, 10*}*, *{*0, 11*}*, and *{*2, 11*}*); and two for P2 and L (*{*6, 10*}* and *{*6, 11*}*).

3. **Generate complete set of states.** [Notations used: *B^n^*, where *n* = 0 to 6, denotes sets of states with *n* number of molecules bound to the enzyme. For example, *B*^0^ is the empty state, and *B*^1^ is the set of one-molecule-bound basis states (see. Figs. 7 & 8). **B***^n^*, for *n >* 1, is the matrix form of each set of bound states, and **B***^n^*, refers to the row of the matrix, where the subscript *m* denotes a state written as a set of basis states (e.g. if *m* = 0, 13, this means state 0, 13). In the matrix, ‘0’ means the states cannot coexist, and ‘1’ means that can coexist. Hence, ‘0’ is equivalent to the ∄ symbol used in Figs. 10 & 11.]

**Fig. 10.**
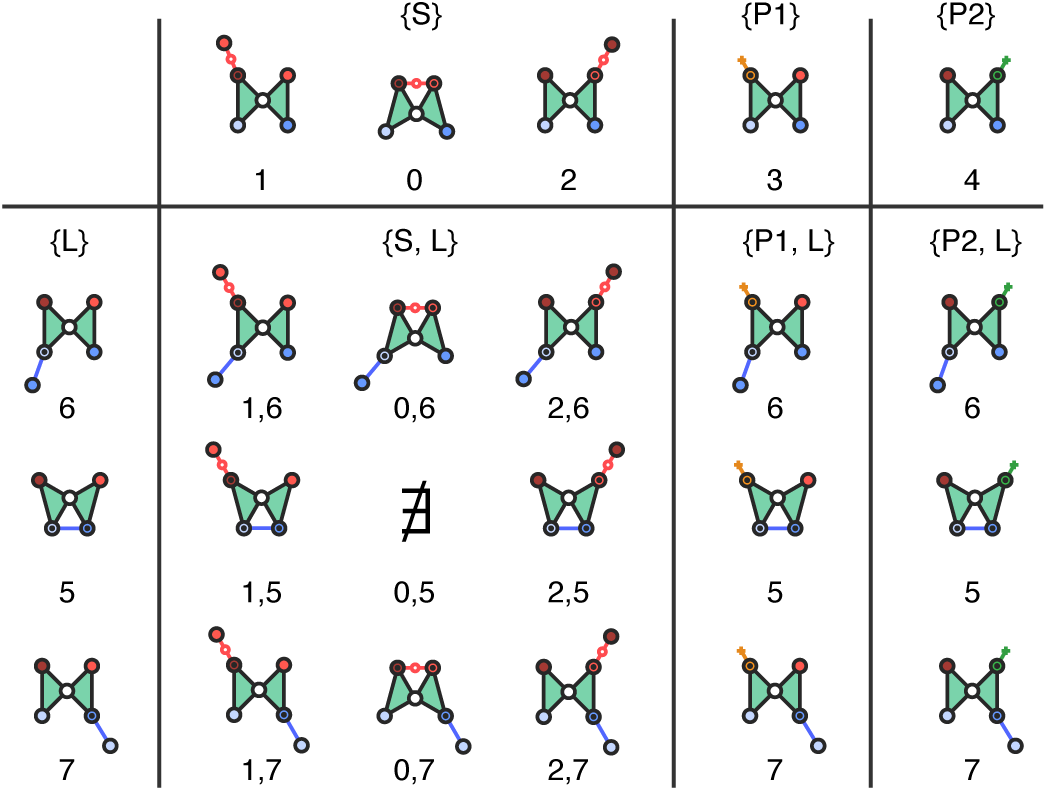
DIV rules matrix. This binary matrix graphically displays the geometric restrictions that exist between the basis states in the DIV system, and which each represent an instance of negative allosteric coupling. The single pairing of basis states that cannot coexist is *{*5*}* and *{*0*}*, or *{*0, 5*}*, which is labeled with ∄, for “does not exist”.

**Fig. 11.**
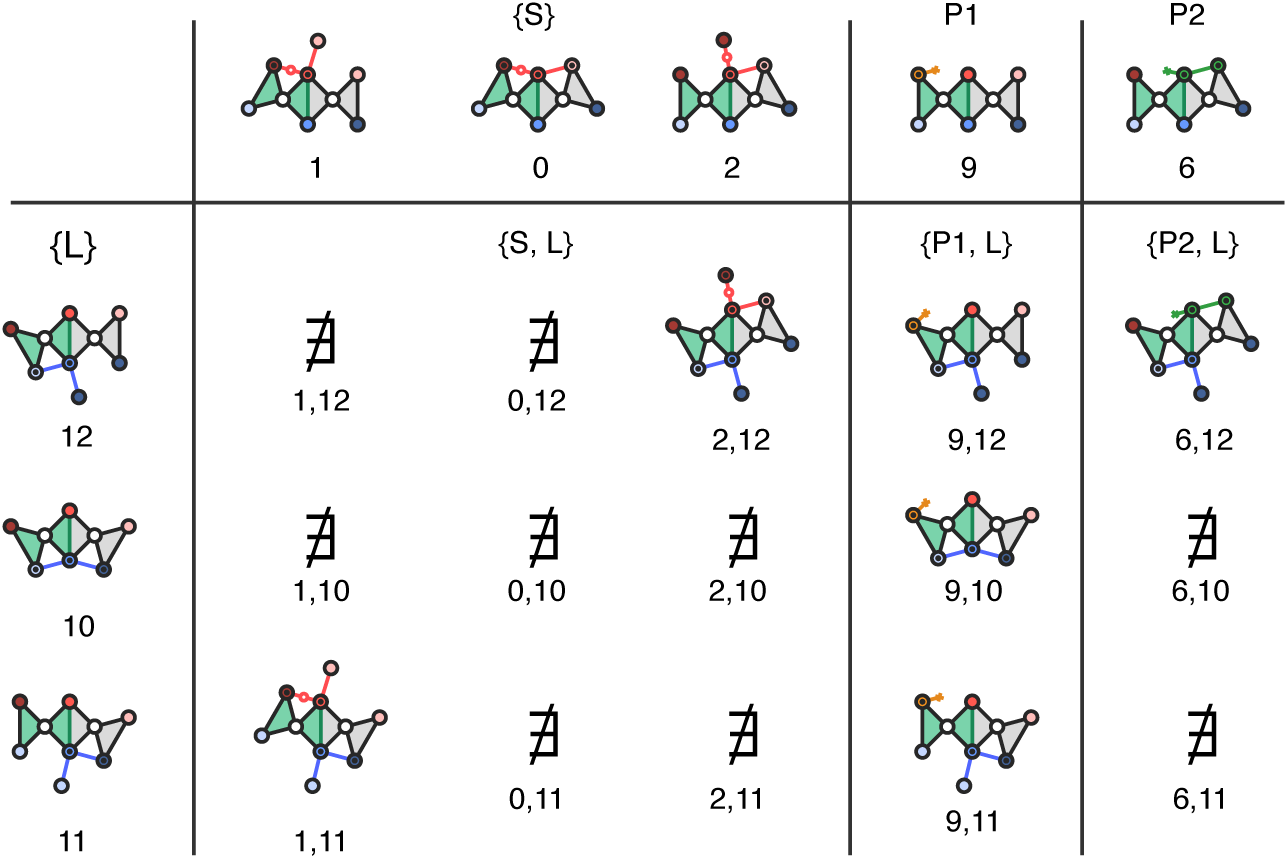
TRV rules matrix. This binary matrix graphically displays the geometric restrictions that exist between the basis states, and which each represent an instance of negative allosteric coupling. Basis states that cannot coexist, and thus generate a new state, are represented with ∄, for “does not exist”. Here, because of the two degrees of freedom, there are nine conflicts, by contrast to the one conflict in the DIV system. Note that P1 and the ligand-bound-states (L) do have any conflicts, because P1 is bound monovalently to the enzyme.

Sets of higher order bound states were successively generated by multiplying (as a binary matrix) the set of states with one molecule less, with the set of basis states *B*^1^ (e.g. *B*^4^ = *B*^3^ *× B*^1^). The process starts with the generation of *B*^2^, where *B*^2^ = *B*^1^ *× B*^1^:

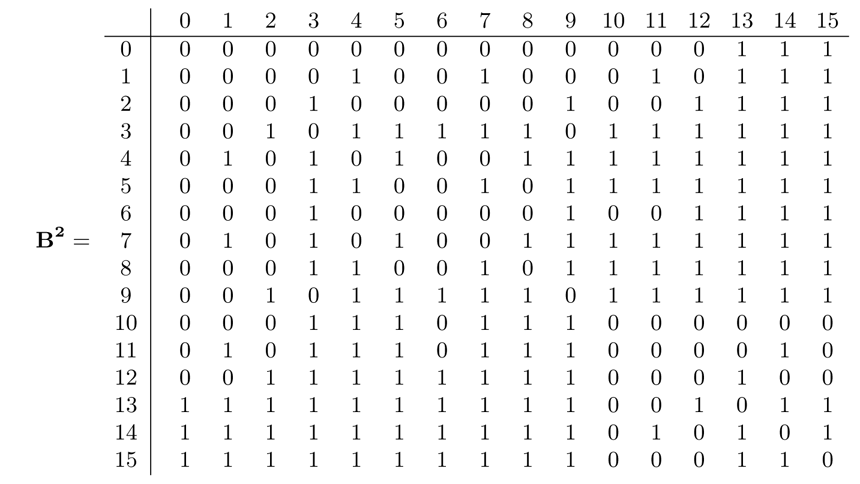

The resulting **B**^2^ matrix has two uses: 1) it generates the two-molecule-bound states, which are indicated wherever a ‘1’ appears in the matrix; and 2) it generates a set of row vectors **B**^2^ to compliment each basis state *m* in *B*^1^, which are in turn used as multipliers to generate the higher order states. For example, the ‘1’ in row 0, column 13, represents state *{*0, 13*}* in *B*^2^. To generate *B*^3^ (*B*^3^ = *B*^2^ *× B*^1^), each state in *B*^2^ is converted into a row vector for **B**^3^, using the set of basis vectors **B**^2^. This is done by performing an element wise multiplication of the basis vectors that comprise each state in *B*^2^. Hence, state *{*0, 13*}* is converted into row vector **B**^3^ in **B**^3^ by multiplying basis vectors **B**^2^ and^2^ together element-wise, which is expressed as the Hadamard product, where ‘*◦*’ means element-wise multiplication:

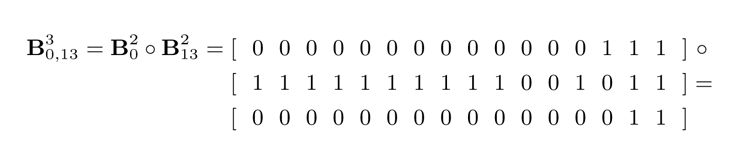

Thus, for every ‘1’ in **B**^2^, the Hadamard product is used to construct the rows of **B**^3^:

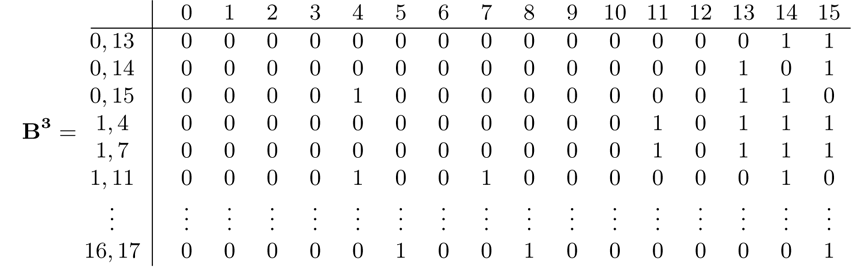

To generate **B**^4^ the process is repeated. For example, the row vector **B**^4^ is generated by the product of **B**^3^ and basis vector **B**^2^ :

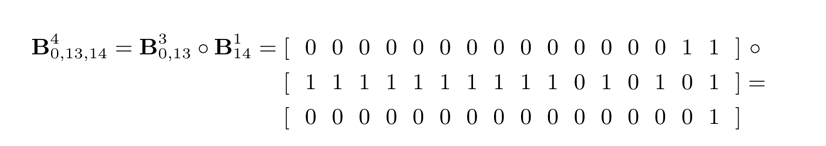

If *BDIV* and *BTRV* are used to symbolize the full set of states in the DIV and TRV systems, respectively, then *BDIV* = *{B*^0^*, B*^1^*, B*^2^*, B*^3^*, B*^4^*}* and *BTRV* = *{B*^0^*, B*^1^*, B*^2^*, B*^3^*, B*^4^*, B*^5^*, B*^6^*}*, where *B*^0^ is the empty state, and the number of higher order sets reflects the number of nodes in each system: four in the DIV system; and six in the TRV system. The number of states in each set is given directly below in Table (). A list of states for the DIV system (*BDIV*) are given in section (), and for the TRV system (*BTRV*) in section 0.3.

4. **Connect states to one another.** The states were connected together in two stages: (i) a unique set was constructed in which each member was a pair of states that could reversibly transition to one another; and (ii) each direction of the pair was again classified by reaction type and assigned with the appropriate rate. The process has some redundancy because pairs were classified by reaction type at both stages. In principle this could have been done once, but this is the way I chose to do it. First I describe stage 1 in more detail, and then stage 1:

**i. Constructing a unique set of transition pairs:** This process was done in three parts: (a) finding pairs that define bimolecular binding/dissociation; (b) finding pairs that define intramolecular binding/dissociation; and (c) finding pairs that define catalysis/ligation. These three categories are described individually below:
a. *Finding bimolecular binding/dissociation pairs:* These pairs were found using a matrix called the binding matrix (**b**), which describes whether two basis states in the *B*^1^ set can combine to form a new state by a binding reaction, with the addition of a row for the empty state (*{*18*}* or *B*^0^):

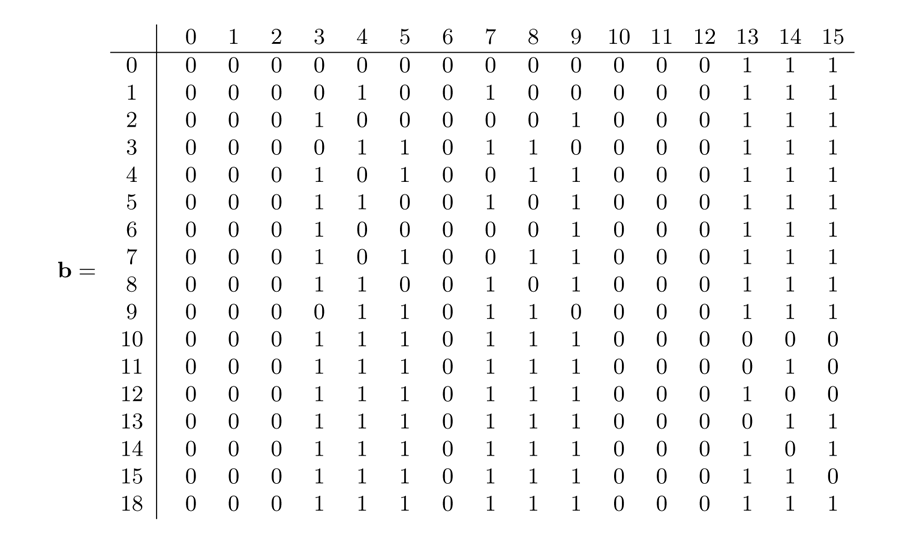

The matrix **b** maps out a subset of the pairs in **B^2^**, with entries that reflect only allowed single-node interactions. An entry of a ‘1’ at a given column position means that the basis state at that column position can combine with the basis state for that row to form a new state that contains the row basis state and the column basis state, where the column state is the new molecule that binds at a single node. Hence, each row can be thought of as a basis vector for binding. And thus, for every state in the system, a combined binding vector could be calculated using the Hadamard product in the exact same way it was done to find successive higher order states. For example, to find the binding vector **b1**,**13** for state *{*1, 13*}*, the binding vectors for *{*1*}* and *{*13*}* are multiplied together:

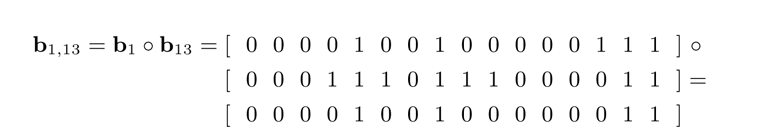

As there is a ‘1’ at positions *{*4*}, {*6*}, {*14*}*, and *{*15*}*, state *{*1, 13*}* can make four different binding transitions to new states containing these basis states, and in reverse, dissociations take place to transition to state *{*1, 13*}*:

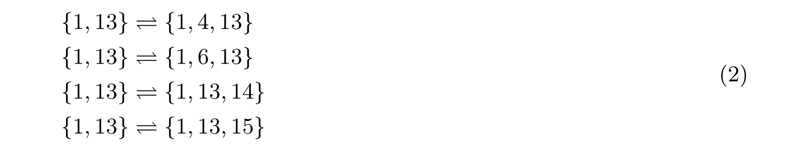

Hence, for every state in the system, the process is repeated to find all the possible bimolecular binding/dissociation transitions.

*b. Finding intramolecular binding/dissociation pairs:* These pairs were found using a matrix, the isomerization matrix (**i**), that describes which states can transition to another by an intramolecular binding/dissociation reaction. For example, when the ligand or substrate are bound at two nodes, and make a third association with the enzyme, and vice versa for dissociation:

Like **b**, **i** is a subset of the pairs defined in **B**^2^. The possible isomerizations indicated by **i** cannot conflict with any other molecule that is bound to the enzyme, where the conflict can be either steric or allosteric. For example, the state *{*1, 14*}* does not have any conflicts, but *{*1, 12*}* does have an allosteric conflict, even though *{*14*}* can transition to *{*12*}* when it is on the enzyme alone. Hence, to account for conflicts, **i** is used with **B**^2^ in the following way. To test for isomerizations, each basis state in a state is tested individually by taking the Hadamard product of its **i** vector and the **B**^2^ basis vectors for all the other basis states in the state. For example, to test state *{*3, 12, 13*}* for isomerizations, *{*3*}*, *{*12*}*, and *{*13*}* are tested separately. Testing *{*3*}* first is done by multiplying **i**3 by **B**^2^ and **B**^2^, which shows that *{*3*}* cannot isomerize when present with *{*12*}*, and *{*13*}*; or substrate on node *a* cannot bind any other node when one ligand is bound divalently at *d* and *e*, and another is bound at *f* :

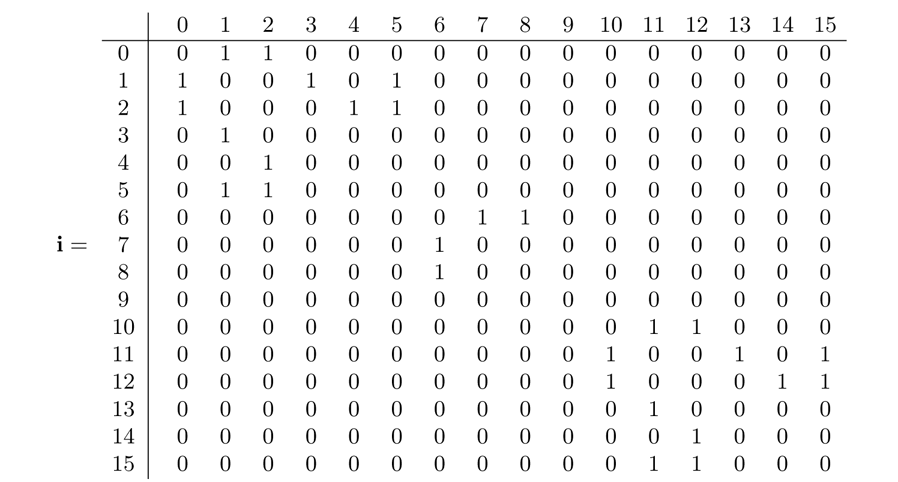

Testing *{*12*}*, reveals one possible isomerization to state *{*3, 13, 14*}*:

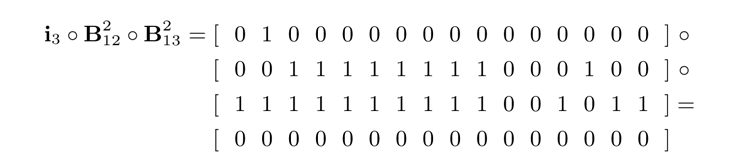

where in this isomerization the divalently bound ligand is dissociating from node *d* or *e*. Testing *{*13*}* results in no other isomerizations. Hence the final result is that *{*3, 12, 13*}* isomerizes to *{*3, 13, 14*}* and *{*3, 13, 15*}*, where because basis states are ordered numerically, *{*14*}* and *{*15*}* are written after *{*13*}* in the new states, though they are isomerization of *{*12*}*:

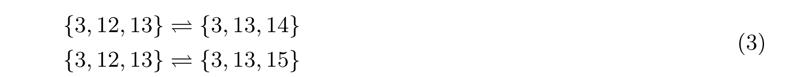

To describe the transitions in terms of the molecules: in the forward direction, the divalently bound ligand dissociates from node *d* or *e*, and in the reverse direction, the same ligand rebinds binds at node *d* or *e*.

*c. Finding cleavage/ligation pairs:* These pairs were found by looking for all states that contain *{*0*}* the fully bound substrate, and converting *{*0*}* to *{*6*}* and *{*9*}*, where *{*6*}* is P1 bound at node *a*, and *{*9*}* is P2 bound divalently. Any state that contains *{*0*}* automatically accounts for the reaction in both directions and rules out any geometric conflicts, because the substrate can only be fully bound when there are no geometric conflicts. Thus any state generated by converting *{*0*}* to *{*6*}* and *{*9*}* is a ligation ready state without geometric conflicts, which specifically means they do not contain basis states *{*11*}* or *{*12*}*.
**ii. Assigning rates to each direction of a reaction pair:** Rates were assigned using a series of logical statements that checked to see what kind of forward and reverse reactions a pair of states described, and then assigned rates accordingly: dissociations were assigned rates based on the node from which the dissociation took place, bimolecular binding reactions were assigned the bimolecular binding rate constant multiplied by a factor that reflected the concentration of the molecule binding, intramolecular binding reactions were all assigned the same unimolecular rate (*k*uni); and cleavage and ligation reactions were assigned the same rate (*k*cat) (see tables 5, 6, and 7, which are repeated from the main text.)

#### 2.4.1 Stochastic formulation of the rates

The rates expressed in table 5 in the SI (table 1 in main text) were input as stochastic rates in the simulations (table 8 in SI). To express the rates stochastically, a scaling factor was used, which is a reaction volume multiplied by Avogadro’s number (*N_A_ v*). By dividing the bimolecular rate constant (*k*_bi_) by this scaling factor, *k*_bi_, which has units of M*^−^*^1^s*^−^*^1^, and which retains some relevance to reaction rates for real biomolecules, was converted into a stochastic version, *k^∗^*, with units of s*^−^*^1^. Likewise, concentrations, *c_m_*’s, which have units of M, were multiplied by the scaling factor to convert them into their stochastic versions, *c^∗^* ’s, which have units of “number of molecules” (No.).

## 2.5 Target cycle of the TRV system with all the transition rates

**Table 1:** Number of states in each set.

| Set | DIV | TRV |
| --- | --- | --- |
| $B^0$ | 1 | 1 |
| $B^1$ | 8 | 16 |
| $B^2$ | 19 | 74 |
| $B^3$ | 17 | 150 |
| $B^4$ | 4 | 142 |
| $B^5$ | - | 58 |
| $B^6$ | - | 8 |
| $\mathcal{B}(\text{total})$ | 49 | 449 |

**Table 2:** Categories of reaction rates.

| Parameter | Value |
| --- | --- |
| Bimolecular rate constant, $k_{\text{bi}}$ | $9 \times 10^6 \text{ M}^{-1} \text{ s}^{-1}$ |
| Concentration, $c_m$ ($m = \text{S, P1, P2, L}$) | varies with $m$; units of M |
| Intermolecular binding rates, $k_m$ | $k_{\text{bi}} \times c_m \text{ s}^{-1}$ |
| Intramolecular binding rate, $k_{\text{uni}}$ | $1 \times 10^6 \text{ s}^{-1}$ |
| Dissociation rates, $k_{\text{off-node}}$ | varies with $node$; units of $\text{s}^{-1}$ |
| Catalysis, $k_{\text{cat}}$ | $100 \text{ s}^{-1}$ |

**Table 3:**
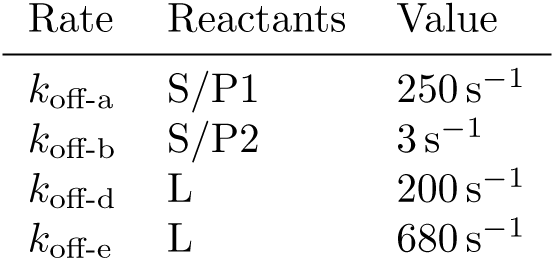
DIV dissociation rates Rate Reactants Value.

**Table 4:** TRV dissociation rates.

| Rate | Reactants | Value |
| --- | --- | --- |
| $k_{\text{off-a}}$ | S/P1 | $250 \text{ s}^{-1}$ |
| $k_{\text{off-b}}$ | S/P2 | $3 \text{ s}^{-1}$ |
| $k_{\text{off-c}}$ | S/P2 | $680 \text{ s}^{-1}$ |
| $k_{\text{off-d}}$ | L | $680 \text{ s}^{-1}$ |
| $k_{\text{off-e}}$ | L | $200 \text{ s}^{-1}$ |
| $k_{\text{off-f}}$ | L | $680 \text{ s}^{-1}$ |

**Table 5:** Stochastic conversions of rates.

| Parameter | Value |
| --- | --- |
| Volume, $v$ | $1 \times 10^{-12} \text{ m}^3$ |
| Scaling factor, $N_A \times v$ | $6.022 \times 10^{11} \text{ M}^{-1}$ |
| Stoch. bimol. rate constant, $k_{\text{bi}}^* = k_{\text{bi}}/(N_A \times v)$ | $9/6.022 \times 10^{-5} \text{ s}^{-1}$ |
| Stoch. “concentration”, $c_m^* = c_m \times (N_A \times v)$ | varies with $m$; units of No. |
| Intermolecular binding rates, $k_m$ | $k_{\text{bi}}^* \times c_m^*$ (or $k_{\text{bi}} \times c_m$) |

## 2.6 Condition for choosing rates

The main condition used for determining the rates was that the energies for the substrate, ligand and P2 satisfied the following inequality:

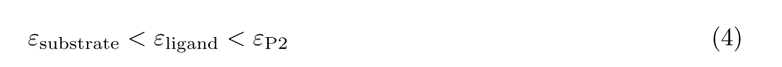

which means that substrate must bind tighter than ligand, which must bind tighter than P2. Each molecule’s energy can be expressed as a summation of its node energies:

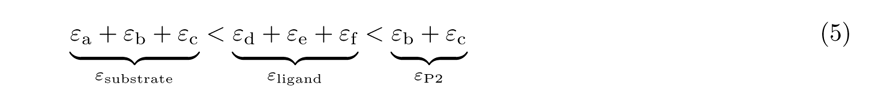

## 2.7 Converting node energies into off rates

Here, an expression is derived for calculating a node dissociation rate (*k*_off-*node*_) in terms of its node energy (*ε_node_*), and vice versa. This is done by satisfying local detailed balance for binding to and dissociation from a single node, as shown in the following figure:

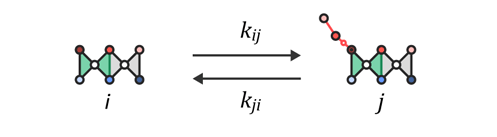

State *i* is the unbound enzyme, and state *j* is the bound enzyme; *k_ij_* is the rate for transitioning from *i* to *j*, and *k_ji_* is the rate for transitioning from *j* to *i*. Although a specific reaction is shown in the figure, the setup of the derivation is general and can work for any node and molecule that binds that node.

If the equilibrium probability of being in state *i* is written as *p_i_*, and for state *j*, written as *p_j_*, then the equilibrium condition must satisfy:

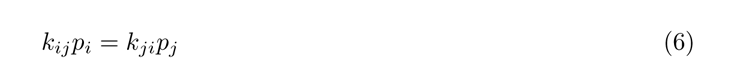

The equilibrium probabilities are given by the Boltzmann factors for each state, divided by the partition function:

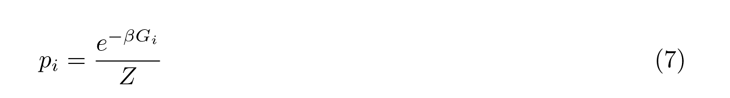

where *G_i_* is the free energy of the state. Using the above expression for *p_i_* and *p_j_* in equation 3, allows the transition rates to be expressed in terms of the state energies:

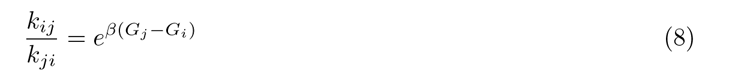

The goal is to replace the above free energies with expressions relevant to states of the linkage. To do this, I closely follow the modeling done in Marzen et al. [71] for a “one-site MWC molecule”, except I leave out the enzyme’s “activated” state and its corresponding energy:

In Table 6, *ε_l_* is the conformational energy of the enzyme (or receptor), and *ε_node_* is the binding energy of the node. The full expression for *µ*, the chemical potential, which is described in Marzen et al. [71] as the “free energy cost of removing a ligand from dilute solution”, is:

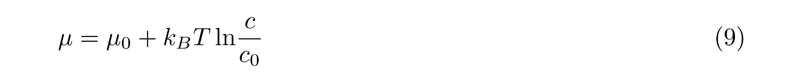

**Table 6.**
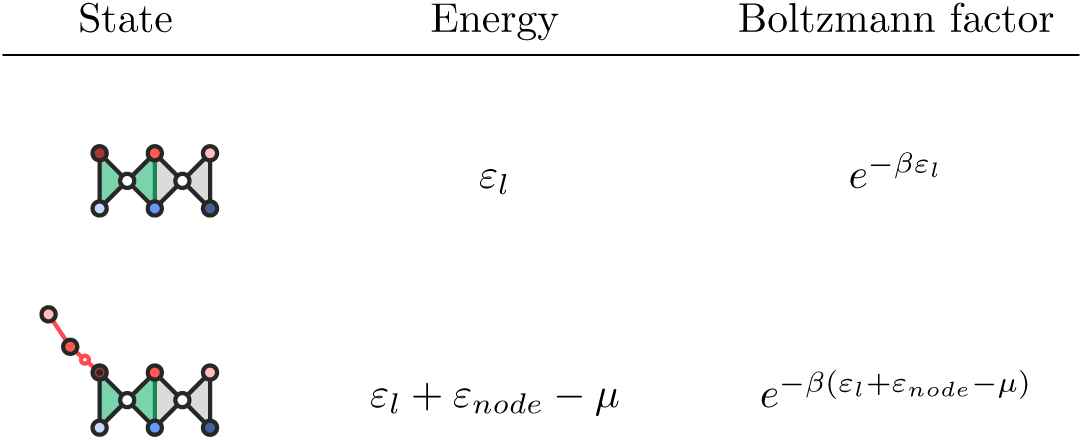

| State | Energy | Boltzmann factor |
| --- | --- | --- |
| | $\varepsilon_l$ | $e^{-\beta\varepsilon_l}$ |
| | $\varepsilon_l + \varepsilon_{node} - \mu$ | $e^{-\beta(\varepsilon_l + \varepsilon_{node} - \mu)}$ |

where *µ*_0_ is an “unspecified reference chemical potential”, and *c*_0_ is an “unspecified reference concentration”. Equation 5 can be re-expressed in terms of the energies in Table 6 by letting *G_i_* = *ε_l_* and *G_j_* = *ε_l_* + *ε_b_ µ*, and by using the full expression for *µ* in equation 6:

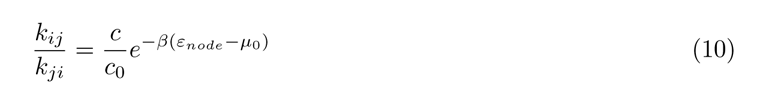

In the final step, *k_ij_*, a binding rate, is expressed as the concentration (*c*) of the molecule binding, multiplied by a bimolecular rate constant (*k*_bi_), so that *k_ij_* = *ck*_bi_; and *k_ij_*, a dissociation rate, is renamed *k*_off-*node*_. Making these two substitutions and solving for *k*_off-*node*_, gives:

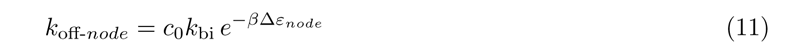

where Δ*ε_node_* = *µ*_0_ − *ε_node_*. Taking the log of both sides allows the node energy to be solved for in terms of the dissociation rate:

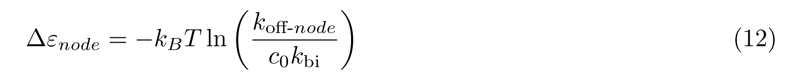

## 2.8 Defining pathways and analyzing trajectories

### Subset of pathways

I define here a subset of thirteen pathways that can be followed on the network displayed in the main text (see Figs. 1I & 2B), using the eight ‘blocks’ of behavior upon which the network is organized. This subset describes the dominant behaviors of the system, and is suitable for describing the simulation results, but it is not general enough to describe the code used to analyze the simulation trajectories. Hence, the full set of pathways is discussed in the next section.

A pathway is defined as a complete fuel cycle, which includes: (1) the loading of substrate; (2) cleavage; and (3) the release of P1 and P2. These steps must take place in the order just given so that the P2 molecular released along that pathway can be attributed to the substrate loaded within that fuel cycle, rather than one in a previous cycle. In addition to this general definition, the subset of pathways considered here obey the following constraints:

i. for non-idling pathways:

a. P1 dissociates before ligand binds
ii. for idling pathways

a. ligand stays bound during idling
b. P1 dissociates before P2

The thirteen pathways are defined using the eight ‘behavior blocks’ (labeled in Figs. 1I & 2B):

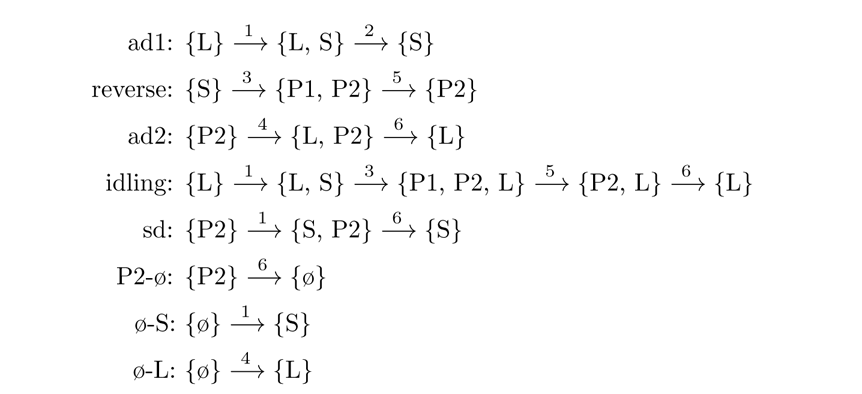

Given the definition of a pathway stated above, idling is both a behavior block and a complete pathway. The remaining twelve pathways constructed from the blocks are:

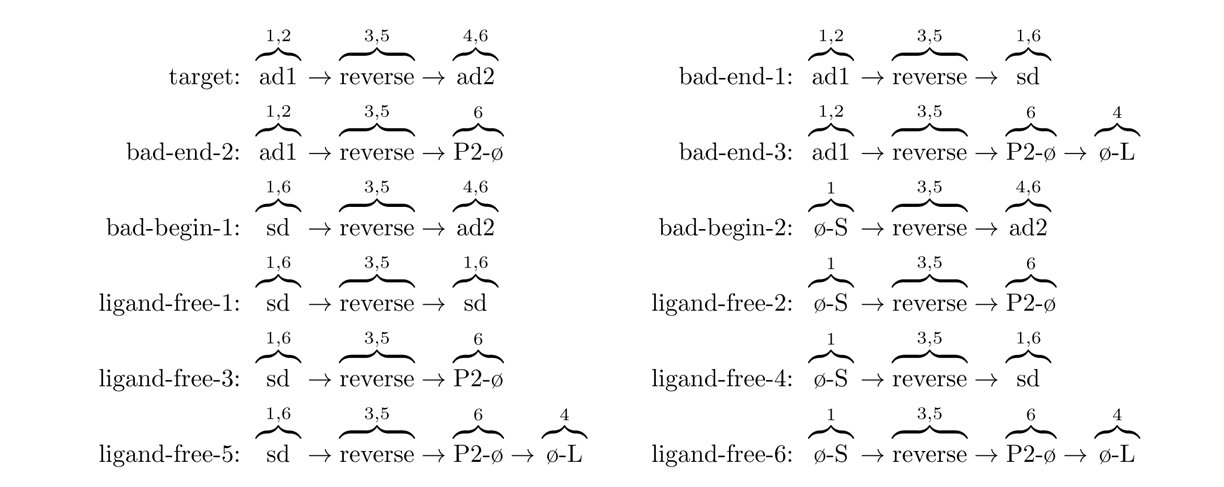

All paths are futile paths, except the first pathway—the target. Each pathway is composed of two halves, which are the two stages of nucleotide exchange: substrate loading and P2 dissociation. The reverse block (cleavage and P1 dissociation) is the crossover point between the two halves. In every futile pathway, one or both stages of nucleotide exchange happen as a futile event (sd, P2-ø, or ø-S). Hence, in the three pathways categorized as “bad-end”, substrate is loaded correctly (ad1), but P2 dissociates through a futile event. In the two pathways categorized as “bad-begin”, substrate is loaded through a futile event, but P2 dissociates correctly (ad2). In the six pathways categorized as “ligand-free”, both substrate is loaded and P2 dissociates through a futile event. And finally in idling, because the ligand is not exchanged, both stages of nucleotide are considered futile.

An important aspect of the ligand-free pathways, as reflected in their name, is that the ligand is not involved in the sequence of events that take place. Thus, the ligand-free pathways reflect the behavior of the system when allosteric coupling has no effect on the operation of the enzyme. Ligand-free-5 and ligand-free-6 are special cases of the ligand-frees in which ligand binds at the very end (see SI 2.8).

### All pathways

In this section I use a kinetic cube to describe the way Python code was written in order to look for the various good and bad pathways that could be taken. The kinetic cube has ten vertices, eight that exist on corners of the cube and two that exist on diagonals. Each of these vertices represents a significant bound state of the enzyme. Pathways drawn on the cube are used to represent changes to the bound state during a simulation.

A complete pathway includes the loading of substrate, catalysis, and the release of P2. To define all the possibilities, pathways were initially broken up into beginnings and endings, which could either be “good” or “bad”, where “good” means it *is* part of a productive cycle, and “bad” means that it is *not* part of a productive cycle. To be a complete productive cycle, the pathway has to begin, and end “good”, though there is only one way to begin “good”. Beginnings define the loading of substrate and the catalysis event, ending with at least P1 and P2 bound to the enzyme, and they start at either L, P2 or ø. Endings define the release of P2 from the enzyme, starting from P1, P2 or P2.

By using a series of logical statements that allowed the beginnings and endings pictured in Fig. 13 to be combined and mutually excluded from one another, all the possible pathways could be enumerated, as visually represented in Fig. 14. Mutual exclusivity here means that the sum of the pathways counted equals the total number of P2’s released during a trajectory. Not every pathway was explicitly searched for. For example, productive cycles 1, 2, and 3 are all possible productive cycles based on the exclusion clauses and the definition of productivity, but they were not differentiated by the Python code that was used.

**Fig. 12.**
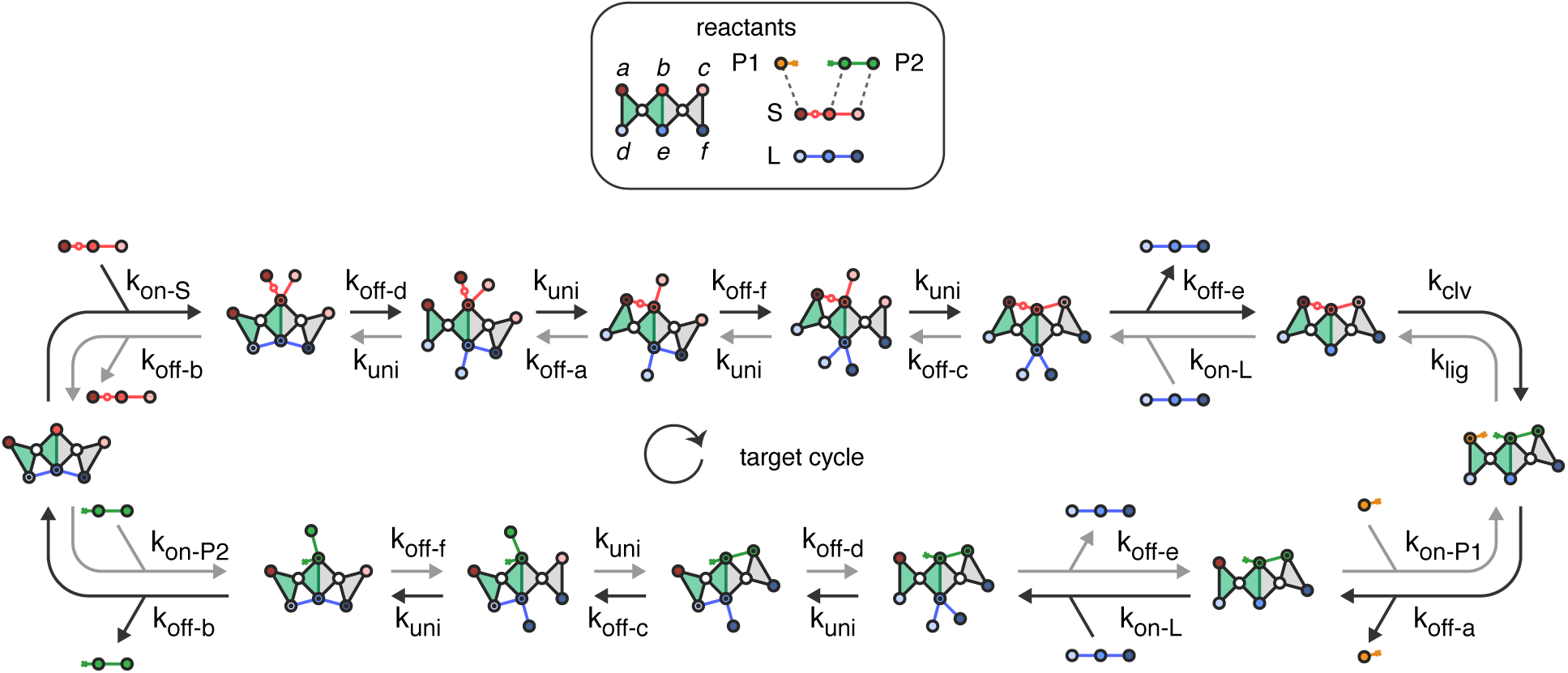
TRV target cycle with rates.

**Fig. 13.**
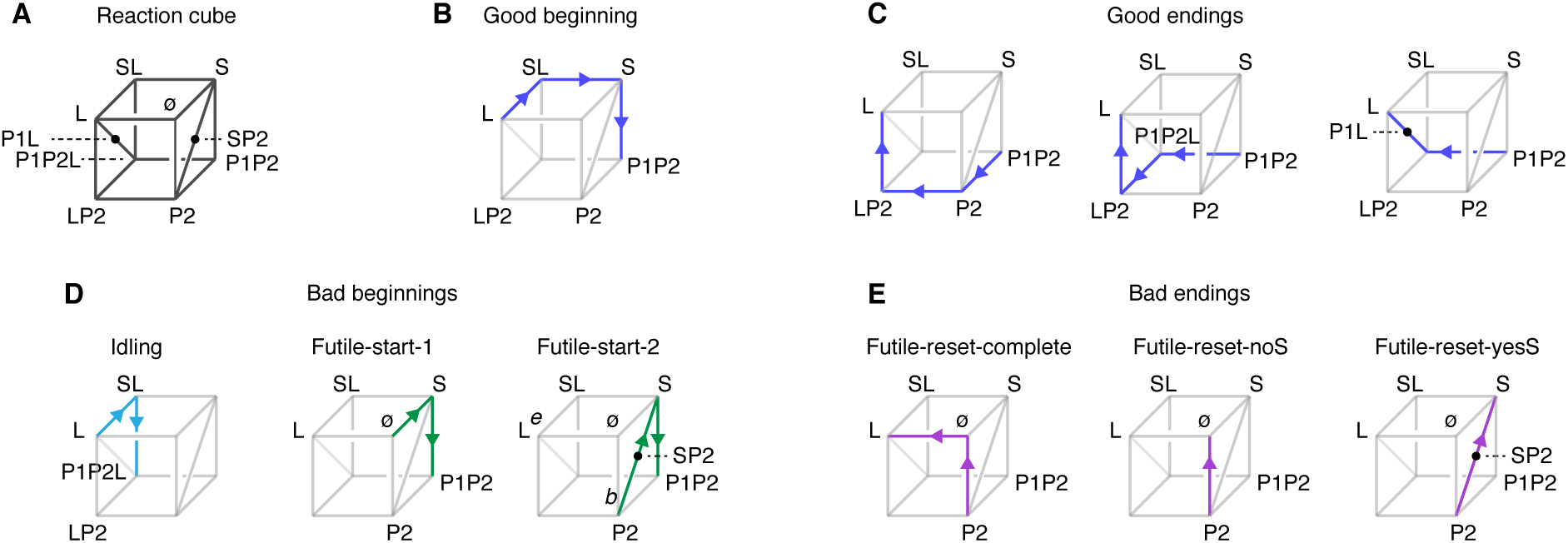
Kinetic cube and partial trajectory pathways. **A,** Kinetic cube. This cube is used to represent possible simulation trajectories. There are ten vertices on the cube - eight on the cube corners and two on the diagonals - which represent the main binding states of the enzyme in combination with S, P1, P2 and L, where the binding state is written without commas or brackets for brevity. **B,** Good beginning. This is the beginning four state sequence of a productive pathway, during which substrate allosterically displaces ligand from the enzyme. **C,** Good endings. These are the three possible ways a productive pathway can end, where in each, ligand binds before P2 dissociates. **D,** Bad beginnings. These are the three bad beginnings, in which substrate is loaded and catalysis takes place, but not in the desired order. In Idling, ligand remains bound during catalysis, and in the futile-starts, substrate is loaded but without ligand being present, thus without the allosteric displacement of substrate by ligand. **E,** Bad endings. These are the three bad-endings, in which P2 dissociates from the enzyme, but without the presence of ligand, thus without the allosteric displacement of P2 by ligand.

**Fig. 14.**
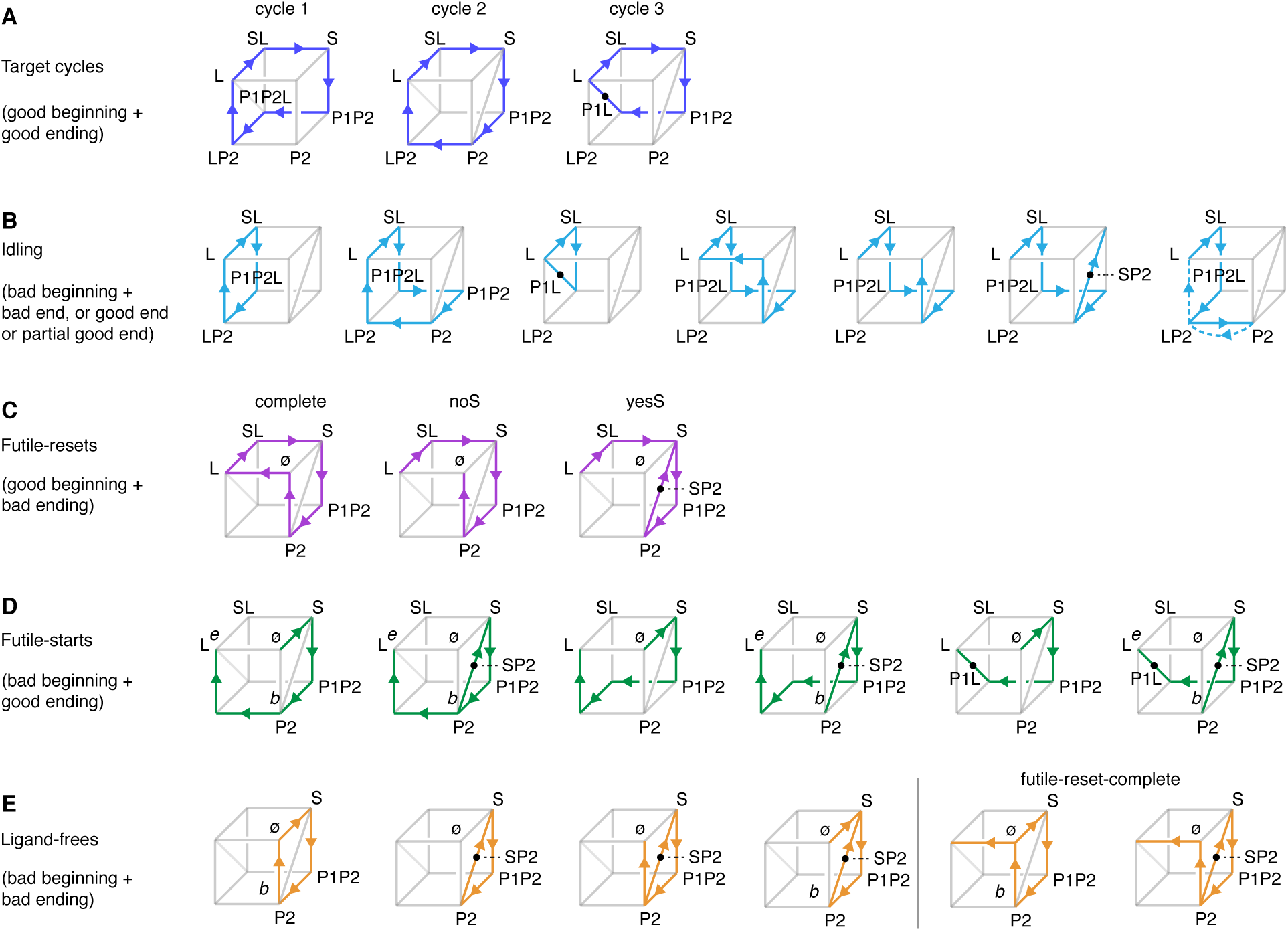
Mutually exclusive pathways on the kinetic cube. Each pathway accounts for one loading of substrate, a catalysis event, and subsequent dissociation of P2. **A,**Target cycles. The three possible target cycles all show the displacement of ligand by substrate, followed by the displacement of P2 by ligand. **B,** Idling. The seven possible idling pathways, all of which start with the ligand remaining bound during catalysis. The first path is the path depicted in Figs. 1I & 2B. In the last pathway, the dotted line segments are the last two steps to take place in the sequence. This is the idling* pathway depicted in Fig. (ref to dwell time plot figure). **C,** Futile-resets. Futile resets start off like a productive pathway - with the displacement of ligand by substrate - but end “off-path” by going through the empty state, or by the steric displacement of P2 by substrate. **D,**Futile-starts. Futile starts begin “bad”, but end “good”, meaning they end with the displacement of P2 by ligand sequence. **E,** Ligand-frees. These are all the pathways that don’t include ligand.

### 2.9 Pathway data

Tables 7 and 8 give the total counts for each pathway. Each total count is a sum of ten counts, where each count is from the ten simulations performed at each of the twenty-four substrate concentrations. Each simulation was run for 100 seconds. Thus, for each simulation (*n*) the turnover rate (*v_n_*), with units of no. P2 released *×* s, is the count for that simulation (*c_n_*) divided by 100 seconds:

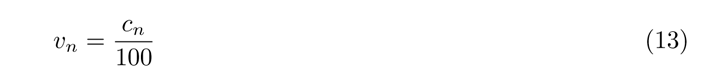

**Table 7:** TRV pathway data. Boxed numbers are the total counts given in Fig. 2F.

| [S] $\mu\text{M}$ | total | idles | be1 | be2 | be3 | bb | lf | lf-frc | ws | pc |
| --- | --- | --- | --- | --- | --- | --- | --- | --- | --- | --- |
| $1 \times 10^{-4}$ | 3 | 0 | 0 | 0 | 0 | 0 | 0 | 0 | 0 | 3 |
| $2 \times 10^{-4}$ | 5 | 2 | 0 | 0 | 0 | 0 | 0 | 0 | 0 | 3 |
| $5 \times 10^{-4}$ | 5 | 2 | 0 | 0 | 0 | 0 | 0 | 0 | 0 | 3 |
| $1 \times 10^{-3}$ | 16 | 5 | 0 | 0 | 0 | 0 | 0 | 0 | 0 | 11 |
| $2 \times 10^{-3}$ | 37 | 9 | 0 | 0 | 0 | 0 | 0 | 0 | 0 | 28 |
| $5 \times 10^{-3}$ | 83 | 20 | 0 | 0 | 0 | 0 | 0 | 0 | 0 | 63 |
| $1 \times 10^{-2}$ | 165 | 36 | 0 | 0 | 2 | 4 | 0 | 0 | 0 | 123 |
| $2 \times 10^{-2}$ | 239 | 59 | 0 | 0 | 0 | 2 | 0 | 0 | 0 | 178 |
| $5 \times 10^{-2}$ | 466 | 132 | 1 | 1 | 1 | 5 | 0 | 0 | 1 | 323 |
| $1 \times 10^{-1}$ | 650 | 169 | 2 | 0 | 0 | 9 | 0 | 0 | 3 | 466 |
| $2 \times 10^{-1}$ | 822 | 220 | 1 | 1 | 0 | 7 | 0 | 0 | 1 | 592 |
| $5 \times 10^{-1}$ | 968 | 239 | 9 | 2 | 0 | 17 | 0 | 0 | 0 | 701 |
| $1 \times 10^0$ | 1051 | 265 | 21 | 3 | 0 | 30 | 2 | 0 | 1 | 728 |
| $2 \times 10^0$ | 1117 | 262 | 53 | 3 | 0 | 55 | 4 | 0 | 1 | 739 |
| $5 \times 10^0$ | 1224 | 324 | 91 | 3 | 0 | 94 | 15 | 0 | 1 | 696 |
| $1 \times 10^1$ | 1269 | 276 | 169 | 3 | 0 | 160 | 39 | 0 | 2 | 620 |
| $2 \times 10^1$ | 1476 | 329 | 221 | 1 | 0 | 208 | 68 | 0 | 6 | 643 |
| $5 \times 10^1$ | 1651 | 339 | 302 | 0 | 0 | 291 | 139 | 0 | 5 | 575 |
| $1 \times 10^2$ | 1666 | 304 | 328 | 1 | 0 | 323 | 179 | 0 | 5 | 525 |
| $2 \times 10^2$ | 1764 | 332 | 334 | 0 | 0 | 329 | 222 | 0 | 6 | 542 |
| $5 \times 10^2$ | 1464 | 269 | 295 | 1 | 0 | 308 | 182 | 0 | 17 | 392 |
| $1 \times 10^3$ | 1028 | 184 | 199 | 0 | 0 | 212 | 153 | 0 | 17 | 263 |
| $2 \times 10^3$ | 441 | 73 | 71 | 1 | 0 | 82 | 71 | 0 | 14 | 129 |
| $5 \times 10^3$ | 103 | 19 | 13 | 0 | 0 | 24 | 19 | 0 | 4 | 24 |

**Table 8:** DIV pathway data. Boxed numbers are the total counts given in Fig. 2G.

| [S] μM | total | idles | be1 | be2 | be3 | bb | lf | lf-frc | ws | pc |
| --- | --- | --- | --- | --- | --- | --- | --- | --- | --- | --- |
| $1 \times 10^{-4}$ | 1 | 0 | 0 | 0 | 0 | 0 | 0 | 0 | 0 | 1 |
| $2 \times 10^{-4}$ | 2 | 1 | 0 | 1 | 0 | 0 | 0 | 0 | 0 | 0 |
| $5 \times 10^{-4}$ | 6 | 2 | 0 | 2 | 0 | 1 | 0 | 0 | 0 | 1 |
| $1 \times 10^{-3}$ | 7 | 5 | 0 | 0 | 1 | 0 | 1 | 0 | 0 | 0 |
| $2 \times 10^{-3}$ | 24 | 4 | 0 | 4 | 4 | 1 | 1 | 2 | 0 | 8 |
| $5 \times 10^{-3}$ | 57 | 16 | 0 | 8 | 18 | 4 | 1 | 4 | 0 | 6 |
| $1 \times 10^{-2}$ | 147 | 35 | 0 | 8 | 41 | 9 | 1 | 17 | 0 | 36 |
| $2 \times 10^{-2}$ | 277 | 61 | 0 | 16 | 75 | 29 | 9 | 34 | 0 | 52 |
| $5 \times 10^{-2}$ | 580 | 123 | 0 | 69 | 128 | 59 | 39 | 73 | 0 | 88 |
| $1 \times 10^{-1}$ | 1010 | 206 | 0 | 150 | 148 | 109 | 110 | 111 | 1 | 175 |
| $2 \times 10^{-1}$ | 1518 | 239 | 4 | 303 | 142 | 209 | 242 | 154 | 1 | 223 |
| $5 \times 10^{-1}$ | 2040 | 311 | 10 | 390 | 90 | 352 | 512 | 94 | 2 | 279 |
| $1 \times 10^0$ | 2477 | 348 | 22 | 486 | 48 | 463 | 750 | 75 | 0 | 285 |
| $2 \times 10^0$ | 2676 | 311 | 37 | 521 | 21 | 529 | 935 | 46 | 0 | 275 |
| $5 \times 10^0$ | 2764 | 372 | 81 | 480 | 12 | 553 | 919 | 16 | 0 | 331 |
| $1 \times 10^1$ | 2753 | 343 | 141 | 411 | 3 | 555 | 990 | 10 | 1 | 299 |
| $2 \times 10^1$ | 2862 | 370 | 235 | 345 | 1 | 592 | 1007 | 2 | 1 | 309 |
| $5 \times 10^1$ | 2888 | 337 | 375 | 193 | 0 | 585 | 1091 | 1 | 3 | 303 |
| $1 \times 10^2$ | 2845 | 364 | 448 | 112 | 0 | 581 | 1003 | 0 | 8 | 329 |
| $2 \times 10^2$ | 2821 | 346 | 450 | 82 | 0 | 547 | 1079 | 0 | 6 | 311 |
| $5 \times 10^2$ | 2722 | 336 | 468 | 37 | 0 | 527 | 1031 | 0 | 18 | 305 |
| $1 \times 10^3$ | 2478 | 290 | 433 | 18 | 0 | 481 | 934 | 0 | 33 | 289 |
| $2 \times 10^3$ | 2233 | 254 | 388 | 11 | 0 | 447 | 834 | 0 | 62 | 238 |
| $5 \times 10^3$ | 1886 | 199 | 292 | 6 | 0 | 378 | 739 | 0 | 107 | 166 |

A mean turnover rate (*v*) was calculated for the ten simulations run:

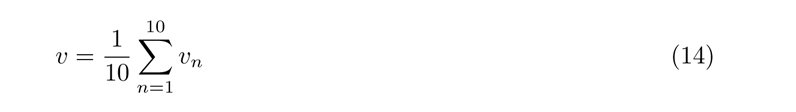

and a standard deviation (sd) of the mean:

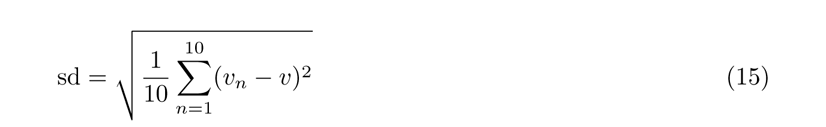

Each point on the plots shows the mean turnover rate for that pathway and its standard deviation.

### 2.10 Allosteric vs steric displacement

Fig. 15, a reaction coordinate plot, illustrates why the allosteric displacement pathway wins over the steric pathway. States along the two pathways are plotted using binding energies that are used for the stochastic simulations. Both pathways need to surmount the same two-part activation barrier of P2 dissociation, to achieve stable binding. This barrier is dissociation of P2 from node *b* (+*ɛ_b_*) and node *c* (+*ɛ_c_*). The difference is that the allosteric pathway can save the highest energy barrier crossing (node *b*) for the end, after ligand has already bound metastably to the enzyme (SI Fig. 15A, states iii and iv). This ordering makes it more probable for the first ligand that binds to remain bound and overcome the barrier. By contrast, multiple binding attempts by substrate are required to overcome the less favorable ordering of the P2 barrier that substrate experiences (node *b*, then node *c*). While both pathways rely on spurious thermal dissociations of P2 for forward progress, these random events can be more easily exploited along the allosteric pathway.

**Fig. 15.**
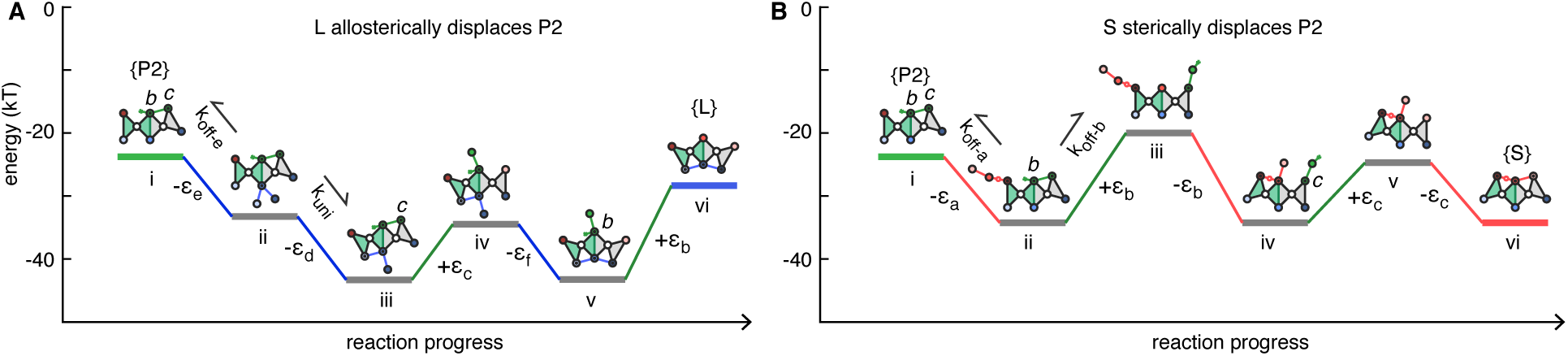
Allosteric displacement vs steric displacement. In this plot the energy landscape traversed by the allosteric and steric displacement pathways are shown on a reaction coordinate, with energy given in kT units, and referenced to the empty state (*{*ø*}*) as zero kT. Both the allosteric and steric displacement reactions must overcome the same uphill activation barrier, which is the energy required to break the two bonds P2 makes with the enzyme (+*ɛ*_P2_ = *ɛ_b_* + *ɛ_c_*)), starting at the P2 state (i).

Along the allosteric pathway (going right from i), the ligand easily overcomes the barrier from the metastable basin it rapidly reaches, after binding divalently to the left side of the enzyme (allosteric displacement; i → iii). From this metastable basin, a single ligand can overcome the two-part barrier in either order, and over many attempts. The figure shows the weaker step being overcome first (iii → v), which is the most probable pathway. Conversely, the steric pathway (going left from i) is more ordered and reversible because the substrate and P2 compete directly for nodes and must alternate. Upon binding, the substrate is initially restricted to node *a* (steric pathway; i → ii), and consequently, the substrate must wait for P2 to dissociate from the more stable *b* node (ii → iii) to make a second bond with the enzyme (iii → iv). This makes it probable for any single substrate that binds to node *a* to dissociate, hence multiple binding attempts by substrate are needed for substrate to sterically displace P2.

### 2.11 Saturation

Saturation (SI Fig. 16) takes place at very high concentrations of substrate (e.g. *>* 200 *µ*M), where the rate at which substrate binds the enzyme surpasses the rate at which bound ligand dissociates from its nodes. At peak saturation, the substrate binding site is occupied by three substrate molecules, and the ligand binding site by one trivalently bound ligand (S, S, S, L)(SI Fig. 16A). The bound substrates cannot displace the ligand because the ligand controls the geometry, and each substrate sterically blocks the other from binding divalently or trivalently. The saturation state persists because if one substrate molecule dissociates, it will likely be replaced by another before the ligand dissociates from one of its nodes and allows a bound substrate to bind divalently.

**Fig. 16.**
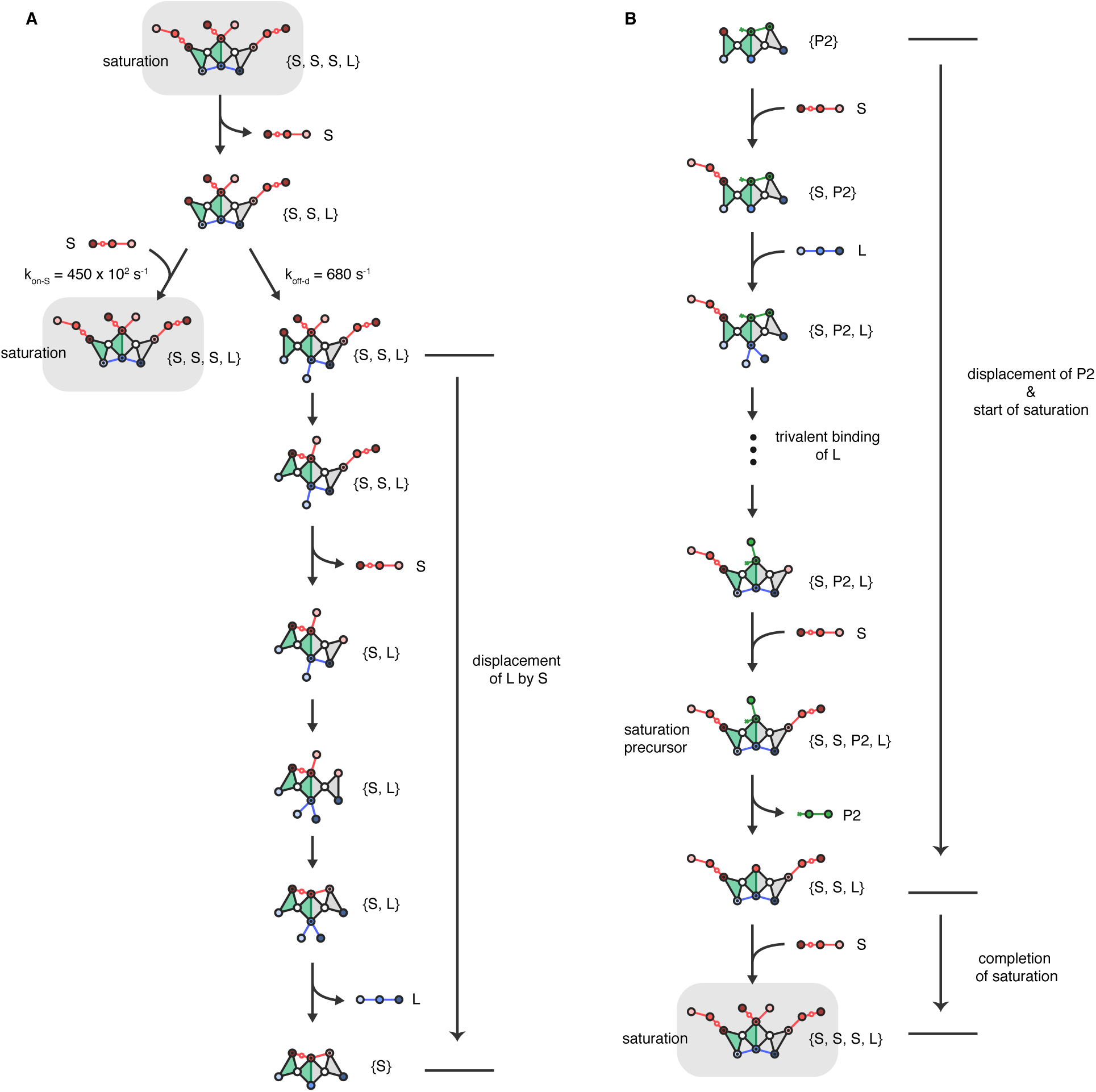
Saturation. **A,** Persistence of the saturation state. The saturation state persists because any substrate that dissociates is likely replaced (left branch) before ligand dissociates from one of its nodes and opens up a pathway for the displacement of ligand by substrate (right branch). Substrate replacement beats ligand dissociation because at high [S], the binding rate surpasses the ligand dissociation by orders of magnitude. For example, at 5 mM [S], *k*_on-S_ = 450 10^2^ s*^−^*^1^, whereas *k*_off-d_ = 680 s*^−^*^1^. The figure shows the saturation state decaying by dissociation of substrate at node a (the second state from top; state 4, 5, 10 in basis form, see SI Fig. 8), but it can also decay by dissociation of substrate from node c (state 3, 5, 10 in basis form). **B,** Pathway to saturation.

The peak saturation state forms during the displacement of P2 by ligand (SI Fig. 16B). During the displacement process, unoccupied nodes at the substrate binding site are rapidly bound by substrate. The first node to be occupied by substrate is a, which becomes available after P1 dissociates. The most likely second node to be occupied by substrate is c, which is a three-step process involving ligand and the displacement of P2 by ligand.

To describe this process, the first thing to note is that spurious dissociates of P2 from c are much more likely than b, because the node c reaction is much weaker than that of node b (*k*_off-c_ = 680 s*^−^*^1^ vs. *k*_off-b_ = 3 s*^−^*^1^). During spurious dissociations of P2 from node c, ligand can bind trivalently by capturing node f and the right degree of freedom, thereby inhibiting P2 from rebinding c (SI Fig. 16B, 4th state from top). In this state, where P2 is still bound at node b and ligand is bound trivalently, substrate will likely bind at node c before P2 dissociates from node b and is released into solution, as the rate at which substrate binds at high [S] is much greater than the rate at which P2 dissociates from b (*k*_on-S_ = 4.5 10^4^ s*^−^*^1^ at [S] = 5 mM^2^ vs. *k*_off-b_ = 3 s*^−^*^1^.). Upon substrate binding to node c, the precursor state of saturation is reached, in which one P2, two substrates, and one ligand are bound to the enzyme (P2, S, S, L)(SI Fig. 16B, 5th state from top). When P2 dissociates from node b, rapid substrate binding to node b will complete formation of the peak saturation complex, in which three substrates and one ligand molecule are bound (S, S, S, L). Peak saturation will dynamically persist, as described above, until one substrate molecule manages to displace ligand.

### 2.12 Maximum turnover rate in the DIV system

The maximum turnover rate (*V*_max_) without ligand is approximately the rate at which it takes the system goes to complete catalysis, P1 dissociation and P2 dissociation (ignoring the substrate binding step which is very fast relative to the other steps at very high concentrations of S):

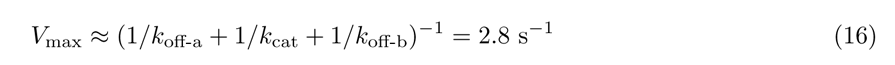

Here, P2 dissociation is rate limiting, because it is the slowest step (*k*_off-b_ = 3 s*^−^*^1^).

### 2.13 Additional plots

In Fig. 17A, I directly compare the ability for ligand to stimulate enzymatic activity in each system, using a measure called the ligand-activation-factor (*A*), which is the ratio of the turnover rate with ligand (*v*_total_), to the turnover rate without ligand(*v*_noL_):

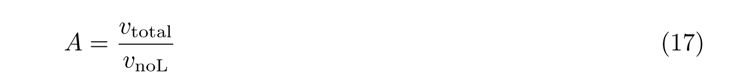

**Fig. 17.**
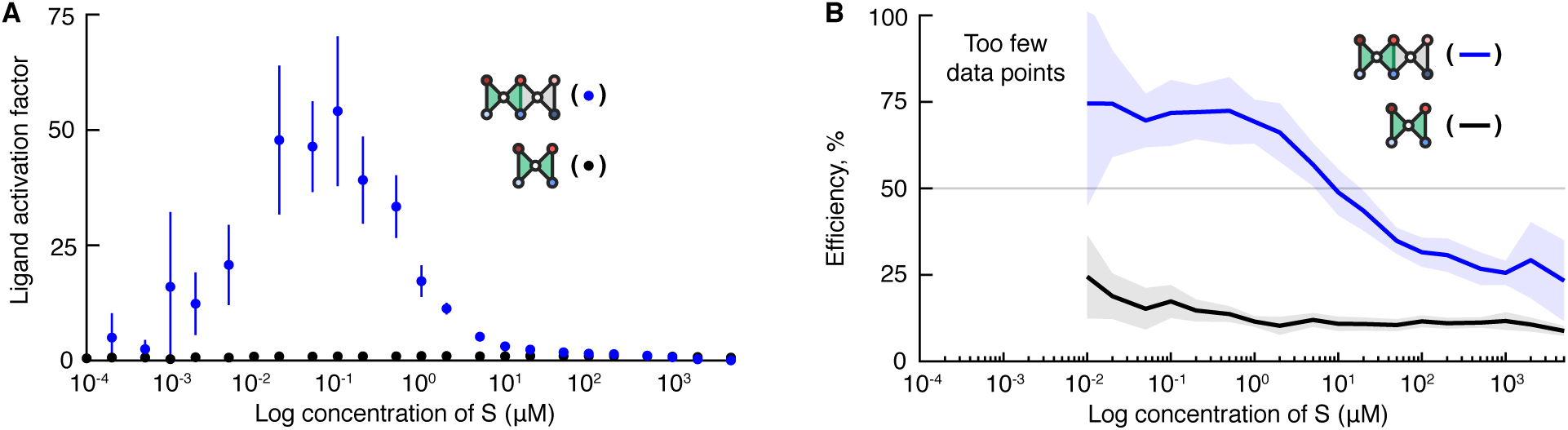
Ligand activation and efficiency. **A,** Ligand activation. Log plot comparing ligand activation (turnover rate with ligand divided by turnover rate without ligand) in the DIV and TRV systems. Error bars are lines around each scatter point. **B,** Efficiency. Log plot comparing efficiencies (target rate *÷* total rate *×* 100) in the DIV and TRV systems.

**Fig. 18.**
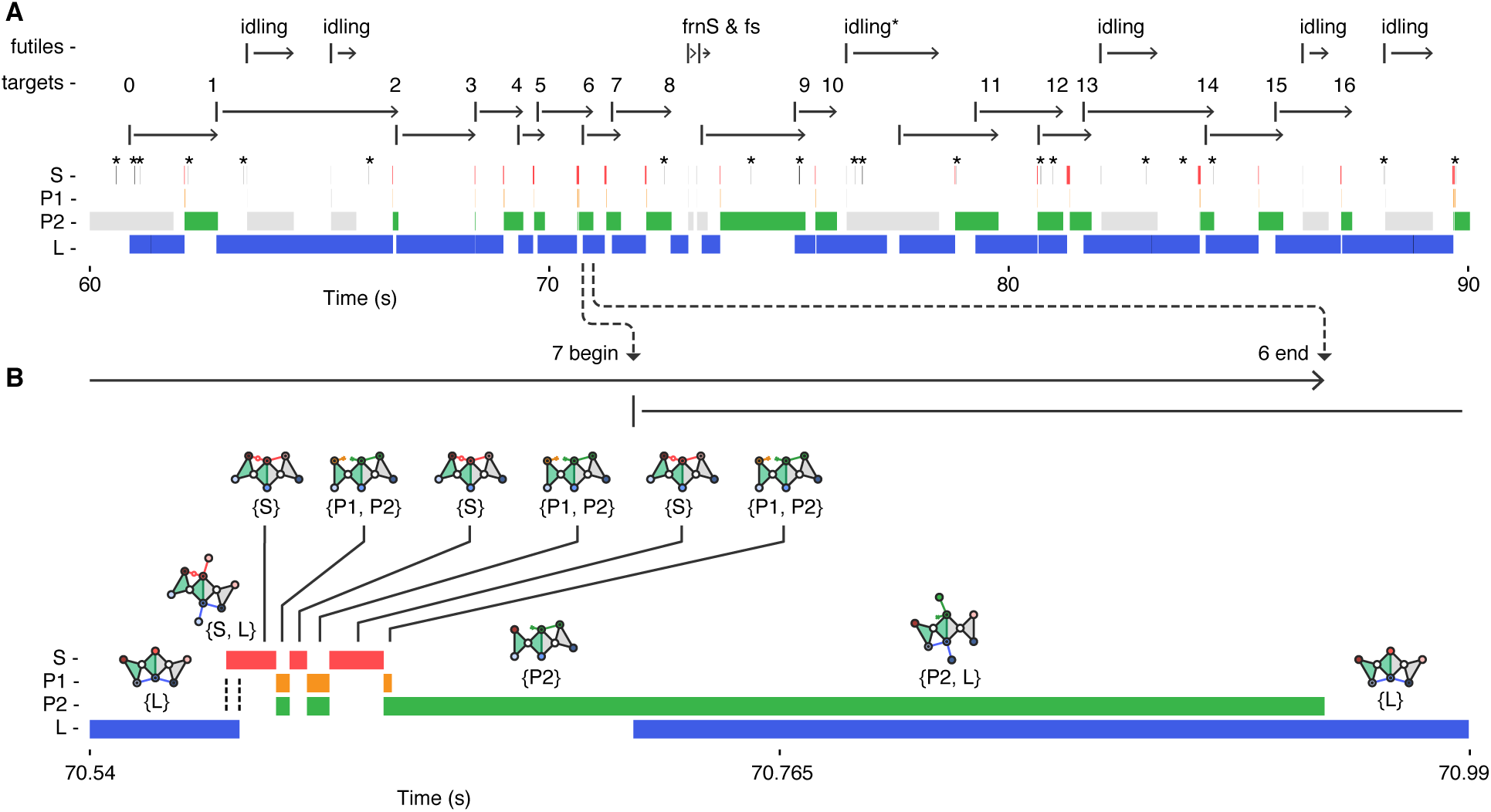
Dwell time plot of a stochastic trajectory, and the reversibility of catalysis. **A,** Thirty seconds of a ninety second trajectory done at 100 nM S and 100 nM L. The time spent bound to the enzyme by each reactant is represented as a horizontal bar of color (S, red; P1, orange; P2, green; and L, blue). Bound states within futile events are colored with grey bars. Spurious substrate and ligand binding events (which are very short lived) are labeled with stars (‘ ’). At the top, the arrows going from left to right span the duration of futile events (idling, frnS and fs) and target cycles (numbered from 1 to 16). Numbers are shown above arrowheads, marking the endings of the target cycles, while vertical bars mark the beginnings. Target cycle arrows are staggered because consecutive target cycles overlap. Note that target cycles start at ligand binding, and end at P2 dissociation - most clearly seen in cycle 11, as it does not overlap with cycle 10. **B,** Reversibility of catalysis. A zoom (70x) of the end portion of target cycle 6 and the beginning of cycle 7, which spans 0.45 seconds of the trajectory. Bound states are represented above by the most connected linkage state in that set. Here, it can be seen how three consecutive cleavage events (red to orange-green overlap) and ligation events (orange-green overlap to red) take place before P1 dissociates (orange-green overlap to green) and rectifies the system. Although not visible in panel **A**, cycles 4 and 14 also display reversible catalysis.

For the DIV system, *A* remains less than one, for all substrate concentrations (Fig. 17A, black dots), though it is difficult to see these small values of *A* in the scale of the plot. Values less than one reflect how the ligand inhibits rather than activates catalysis in the DIV system. By contrast, in the TRV system, *A* rises to over 50x at [S] = 0.1 *µ*M (Fig. 17A, blue dots). The ability of the ligand to act as a switch in the TRV system, by turning on enzymatic activity for a range of substrate concentrations, or likewise, for the enzyme to be relatively dormant in the presence of substrate alone, reproduces actin-activated ATPase activity in myosin, in which myosin processes ATP up to 700x times faster in the presence of actin [19]. The mechanism is the same too - tightly bound product must be “exhausted” though an allosteric displacement, which empties the reactive site so the next cycle can begin.

In Fig. 17B, I directly compare efficiencies of the two systems. The efficiency (*E*) is defined as the percentage of P2’s made and released by the target cycle (*v*_target_), with respect to the total turnover rate with ligand (*v*_total_):

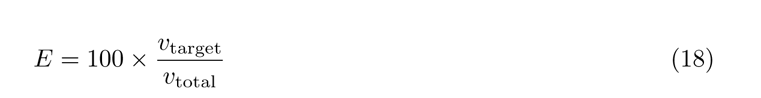

SI Fig. 17B shows that the TRV system maintains an efficiency close to seventy percent for concentrations of substrate between 0.02 *µ*M and 1 *µ*M, and then falls below fifty percent above 10 *µ*M [S]. By contrast, the efficiency of the DIV system does not rise above twenty-five percent.

In an attempt to visually capture the cyclic and also stochastic behavior of the TRV system, I plot thirty seconds of a simulation trajectory, where the time spent bound to the enzyme by each of the four reactants (a dwell time) is represented as a horizontal bar of color (SI Fig. 18). The figure is useful for visually depicting single-enzyme behavior that cannot be seen in the turnover plots - behaviors relevant to understanding the operation of the TRV system.

One behavior visible in SI Fig. 18 is that the enzyme spends all its time bound to at least one molecule, with only one exception between cycles 8 and 9, where a bad-end-2 is followed by a bad-begin-1. Overlap between consecutive target cycles, which takes place when ligand displaces P2, is also visible (see cycles 11 to 13 for example). Most of the target cycles overlap with their preceding and trailing target cycles, except for cycles 1, 9, and 11.

Another behavior visible in SI Fig. 18 is how idling can be interpreted as a “stumble”, rather than complete “trip” to the system. All incidences of idling, except one, are followed by a target cycle, where the ligand bound at the beginning of the target cycle is the same ligand bound throughout the idling event. In other words, these idling events can be regarded as taking place “within” the target cycle that follows them. For example, the first two idling events which take place can both be regarded as taking place within target cycle 2. The exception takes place between the end of target cycle 10 and beginning of 11, where the ligand molecule bound dissociates during the idling event (SI Fig. 18, see idling*).

In general, SI Fig. 18 allows a visual comparison of dwell times in the target cycle. The enzyme spends most of its time bound to ligand, P2 or both, and a relatively little amount of time bound to substrate. The difference means that allosteric displacement of ligand by substrate (overlap of red and blue) is much more rapid than allosteric displacement of P2 by ligand (overlap of green and blue). A single instance of this comparison is easily seen in SI Fig. 18B, which is a 70x zoom of target cycle 6. The dwell time pattern is consistent, though a systematic analysis was not carried out in this study.

The most mechanistically significant behavior visible in SI Fig. 18B is the reversibility of the catalysis reaction. In the zoomed in view of cycle 6, the cleavage reaction (S → P1, P2 ; red to orange-green overlap) is twice reversed by a ligation reaction (P1, P2 → S ; orange-green overlap to red), until P1 dissociates after the third cleavage reaction to rectify the reaction. Although not magnified and thus not visible, cycles 4 and 14 also display reversible catalysis. These cycles, which ultimately go forward despite reversing at the catalysis stage, reflect how the direction of the catalysis reaction is controlled by the respective stabilities of the species bound at either end of the cleavage and ligation reactions (S vs P1 and P2), rather than the relative rates of cleavage vs ligation, which are set equal to one another.

### 2.14 Mapping to Purcell’s three-link swimmer

Mechanical cycle I, which goes through *C*_2*A*_, is non-reciprocal in the conformational transitions it makes, whereas mechanical cycle II, which goes through *C*_2*B*_, is reciprocal. To show this difference, the linkage enzyme is mapped to Purcell’s three-link swimmer [30]. The three-link swimmer is a chain of three bars connected at two flexible joints (SI Fig. 19B). Motion of the two outside bars, around each of their joints, is captured by two angle parameters, *θ*_1_ and *θ*_2_. In phase space, cycle I produces a closed curve, and cycle II produces an open curve (SI Figs. 19C & 19D). Mechanical cycle I is non-reciprocal because it breaks time-reversal symmetry—the sequence of conformational changes in forward time is different from the sequence in reverse time (SI Fig. 19E). By contrast, the sequence of conformational changes in mechanical cycle ii are reciprocal because they are the same in forward and reverse time (SI Fig. 19F).

**Fig. 19.**
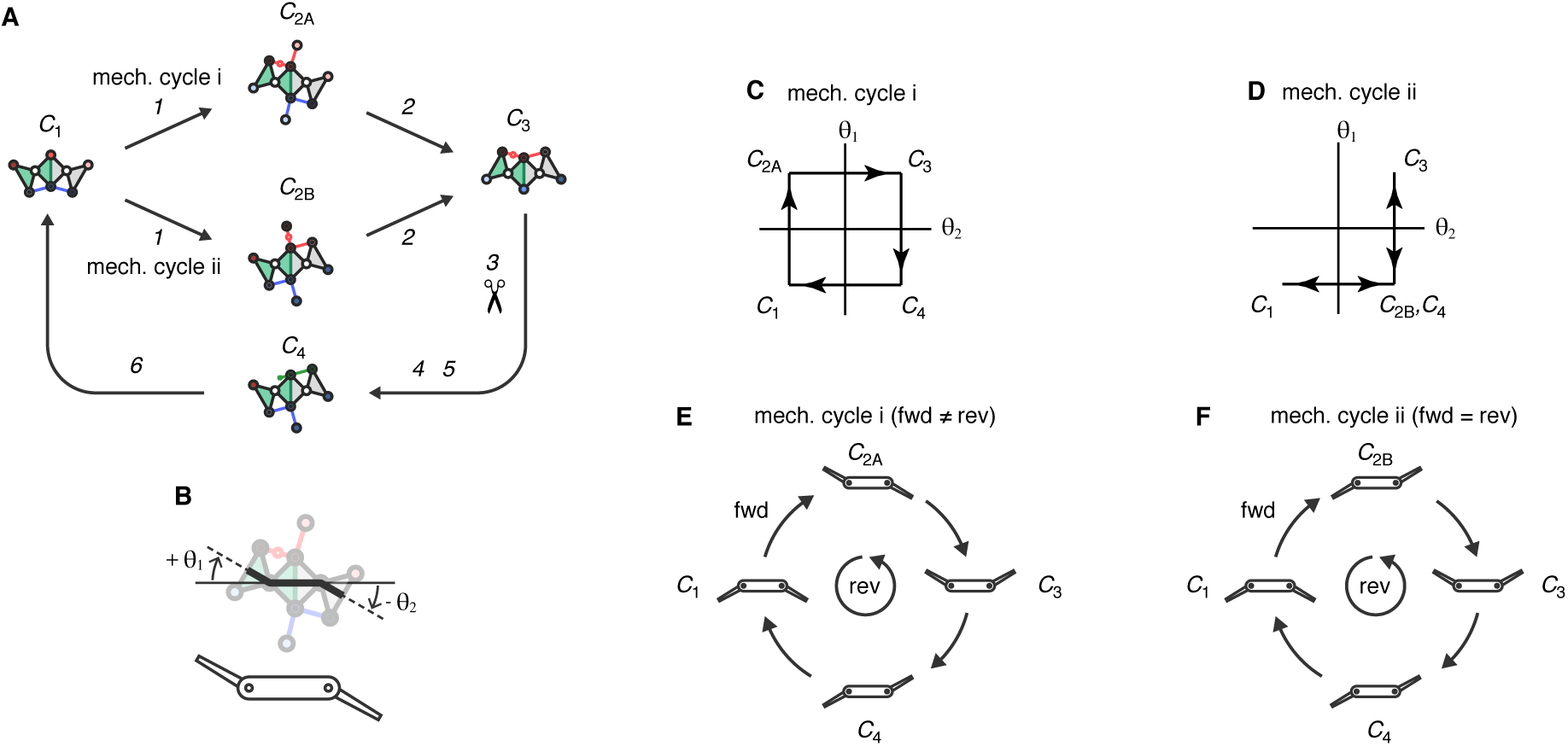
Two mechanical cycles of the 2unit linkage and their mapping to Purcell’s. [30] **three-link swimmer**. **A,** Mechanical cycles. The cycle displayed (left) shows both mechanical cycles, each of which contain four rigid states, labeled *C*_1_ − *C*_4_ (*C* for ‘conformation’), where the P1, P2 state is included after *C*_3_ because it is the point at which flexibility is introduced into the cycle, which allows both chemical and mechanical reversal takes place. The degeneracy that leads to two mechanical cycles, which both go through the same chemical cycle (right), takes place at *C*_2_ (S, L). The complete sequence for cycle I is shown in Fig. 1H (See SI for a complete cycle ii). **B,** Three-link swimmer mapping. The linkage can be mapped to a three-link system, where two outside bars move relative to a ‘stationary’ center bar, and their up and down states are reported by *θ*_1_ and *θ*_2_. **C,** Mechanical cycle I. In the phase space (*θ*_1_*, θ*_2_), mechanical cycle I produces a closed curve. **D,** Mechanical cycle ii. By contrast, mechanical cycle II produces an open curve. **E,** Time-reversal asymmetry of mechanical cycle I. The forward and reverse time conformational changes of mechanical cycle I are not equivalent (*C*_1_ *→ C*_2_ *→ C*_3_ *→ C*_4_ =*̸ C*_1_ *→ C*_4_ *→ C*_3_ *→ C*_2_). **F,** Time-reversal symmetry of mechanical cycle II. The forward and reverse time conformational changes of mechanical cycle II are equivalent (*C*_1_ *→ C*_2_ *→ C*_3_ *→ C*_4_ = *C*_1_ *→ C*_4_ *→ C*_3_ *→ C*_2_).

**Fig. 20.**
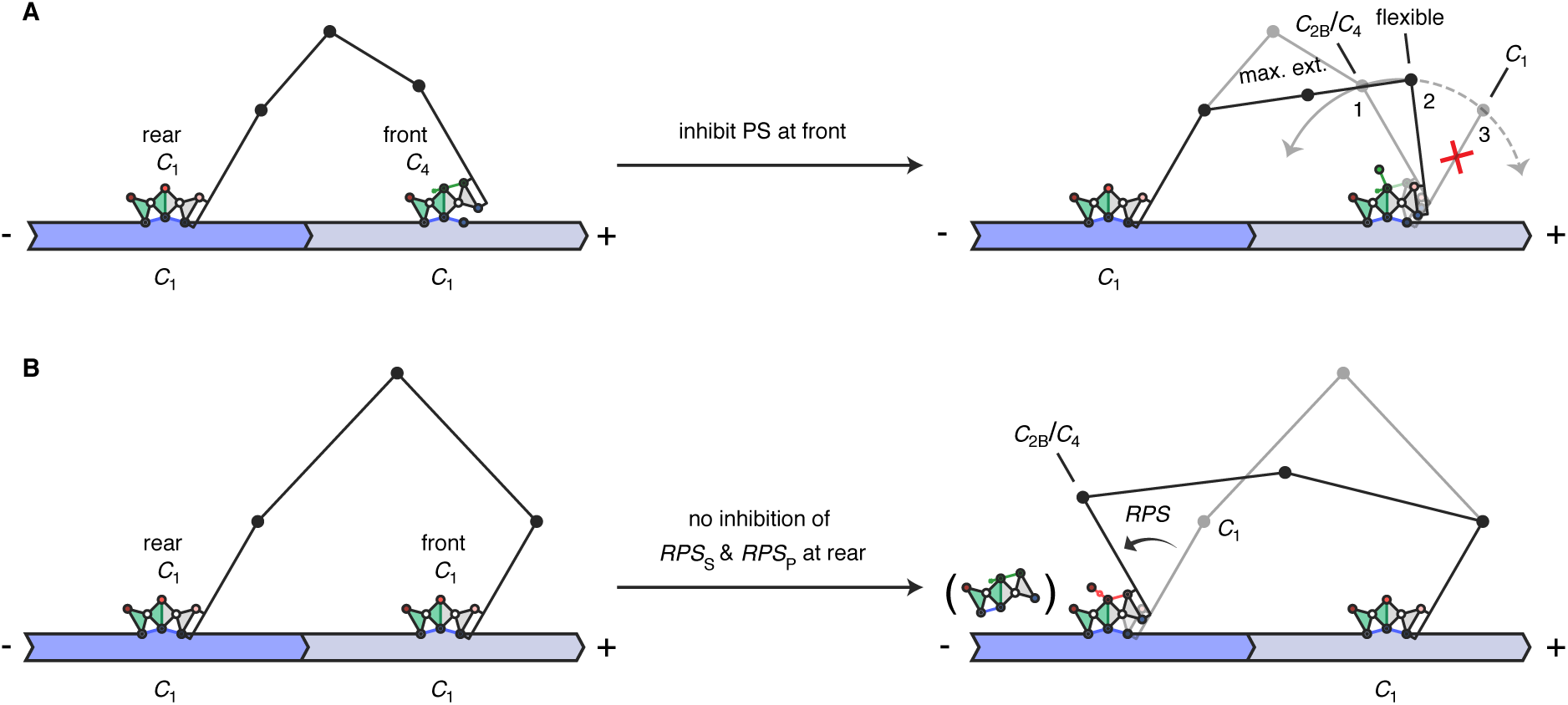
Pick-up bias walker and no-bias walker. **A,** Pick-up bias walker. The power stroke (PS) is inhibited in the front unit by allosteric feedback. **B,**

**Fig. 21.**
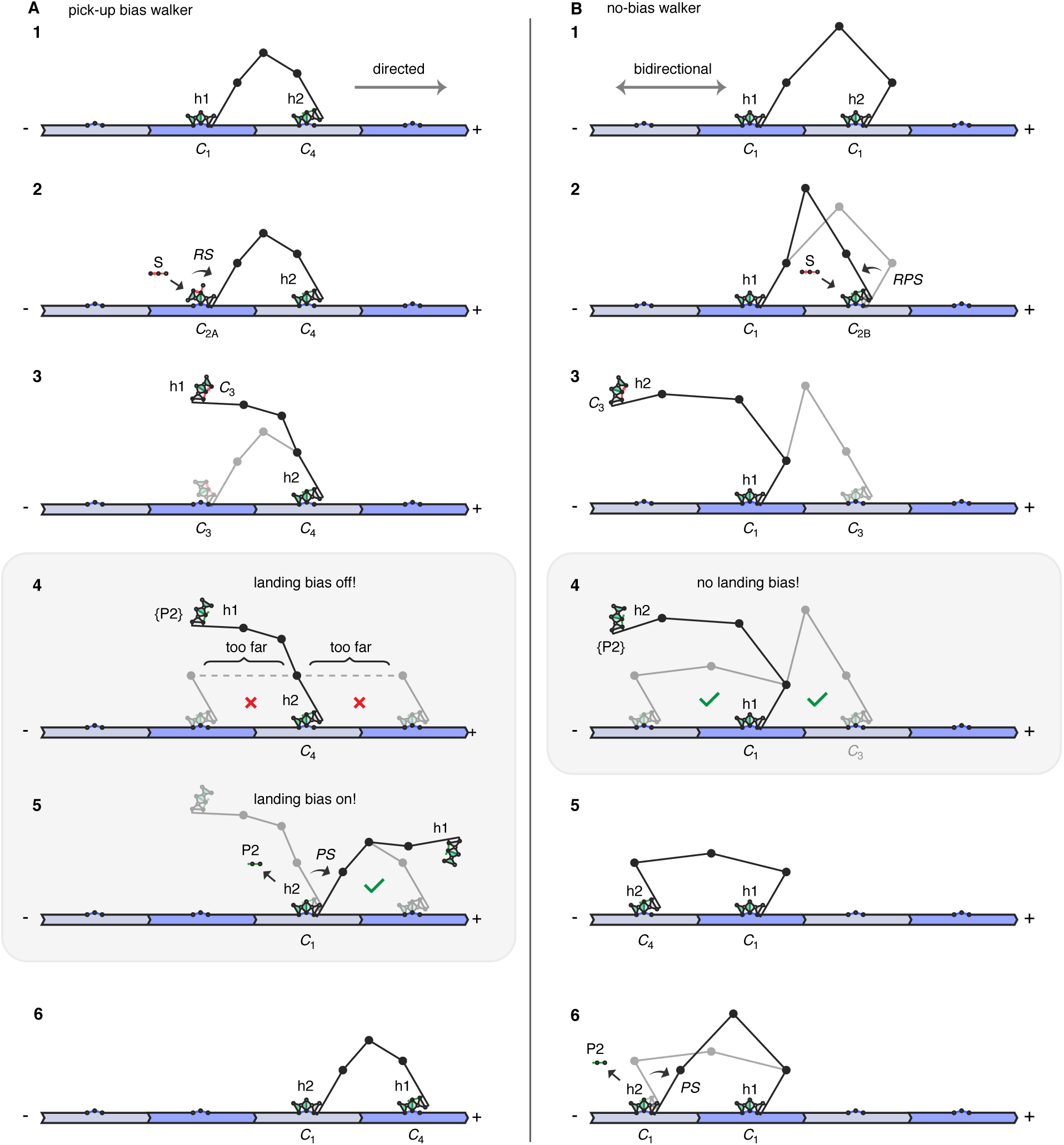
Stepping in the pick-up and no-bias walkers. **A,** Forward step of the pick-up bias walker. **B,** Backwards step of the no-bias walker.

### 2.15 Pick-up bias walker and no-bias walker

### 2.16 States of the DIV system

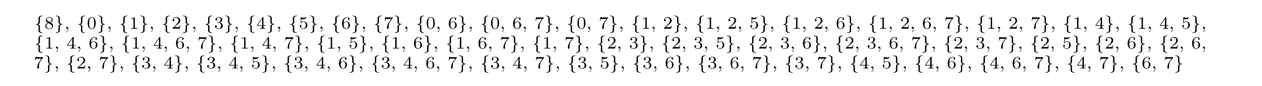

### 2.17 Transitions out of each state in the DIV system

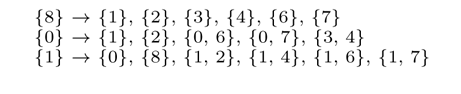

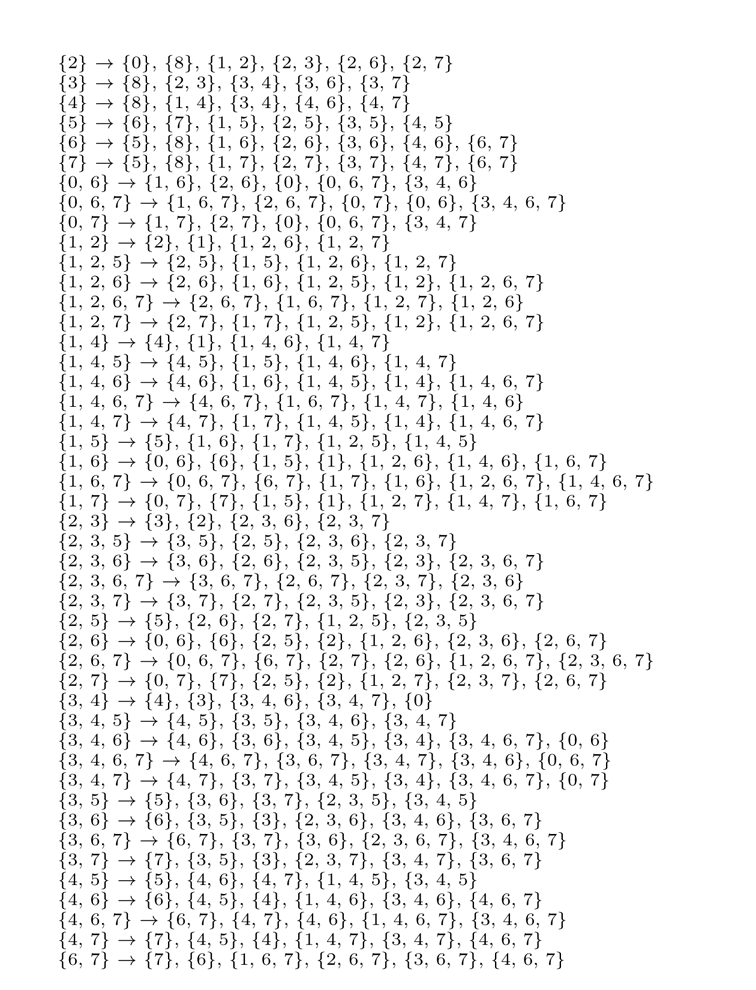

### 2.18 States of the TRV system

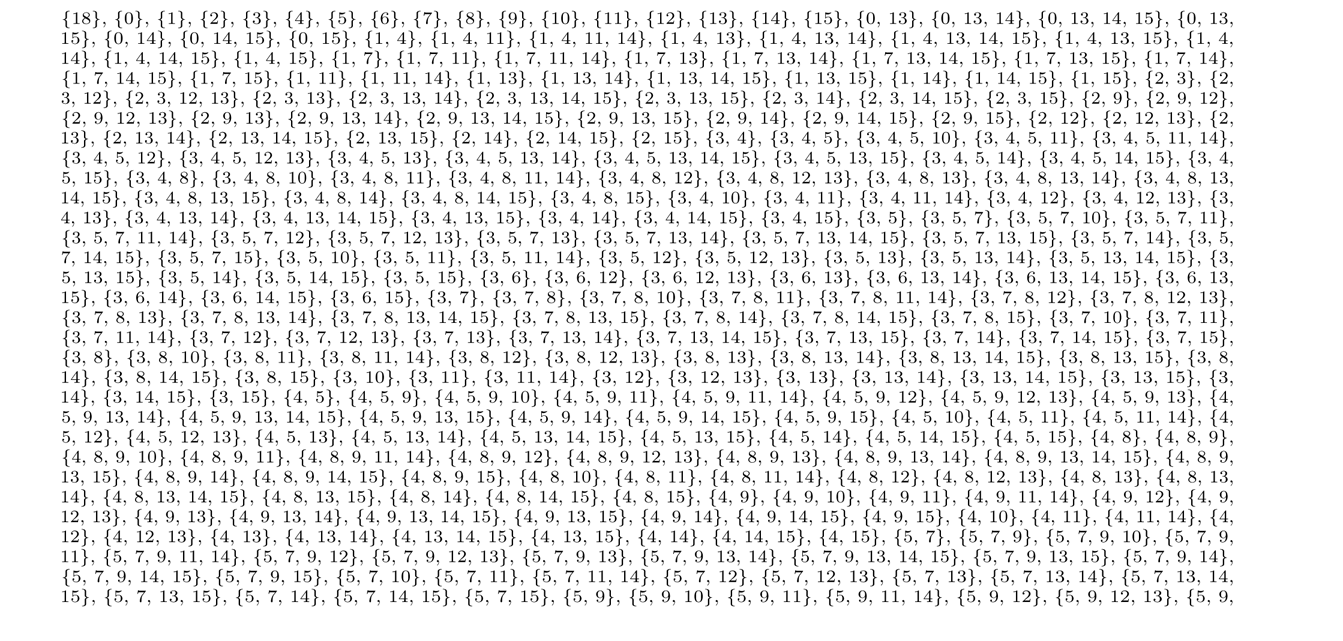

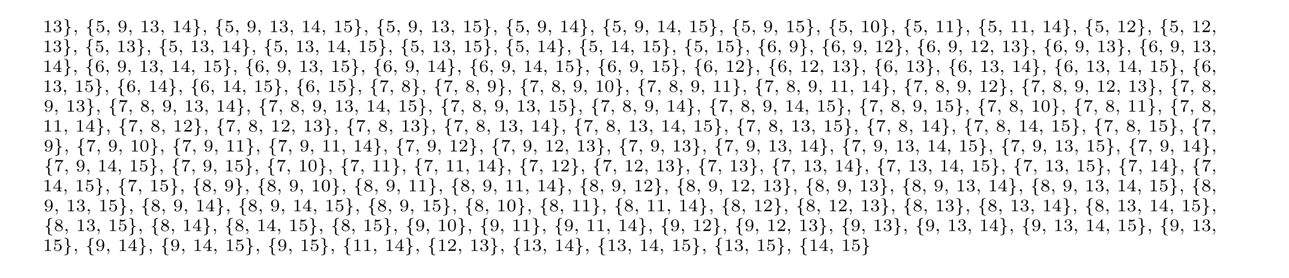

### 2.19 Transitions out of each state in the TRV system

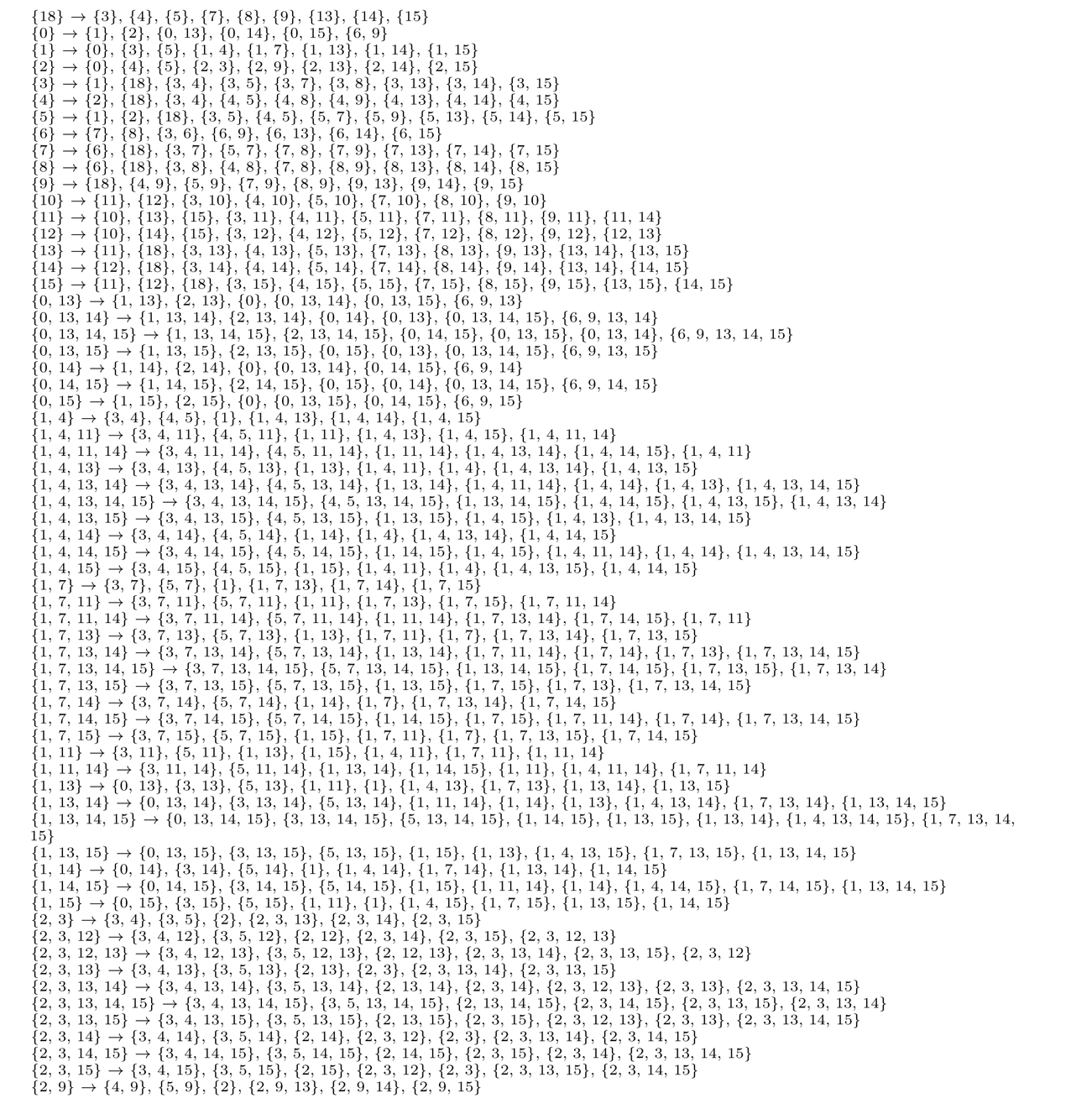

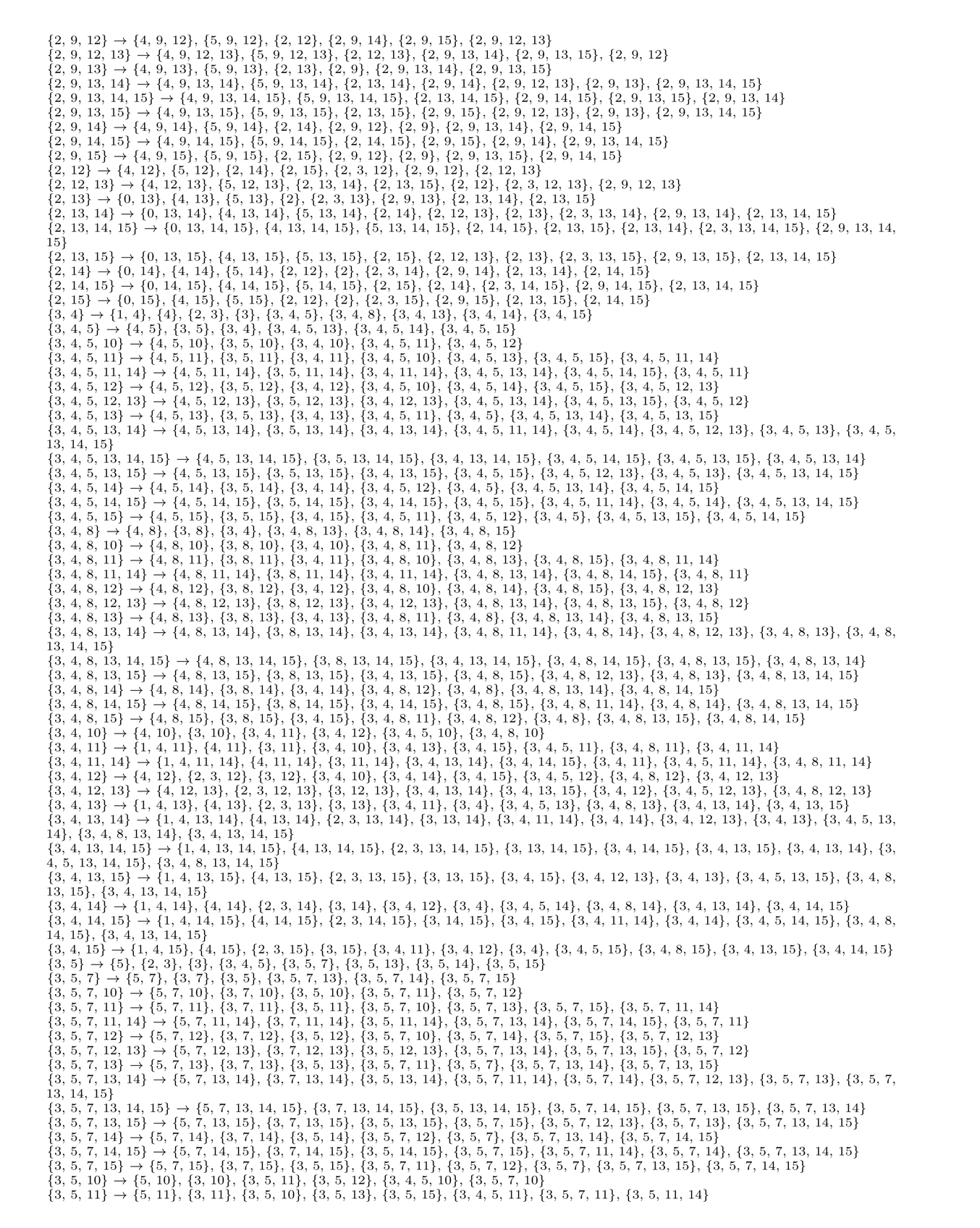

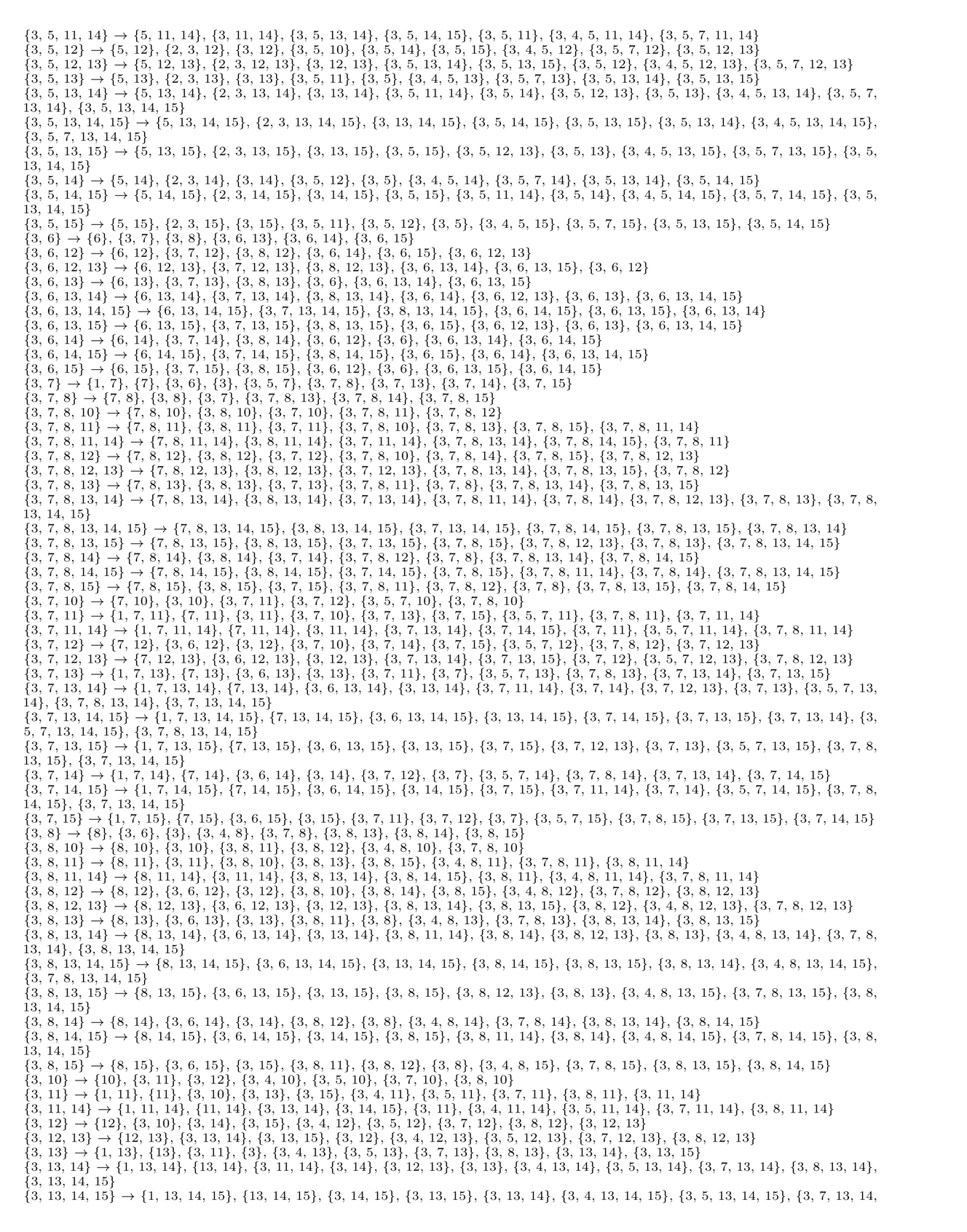

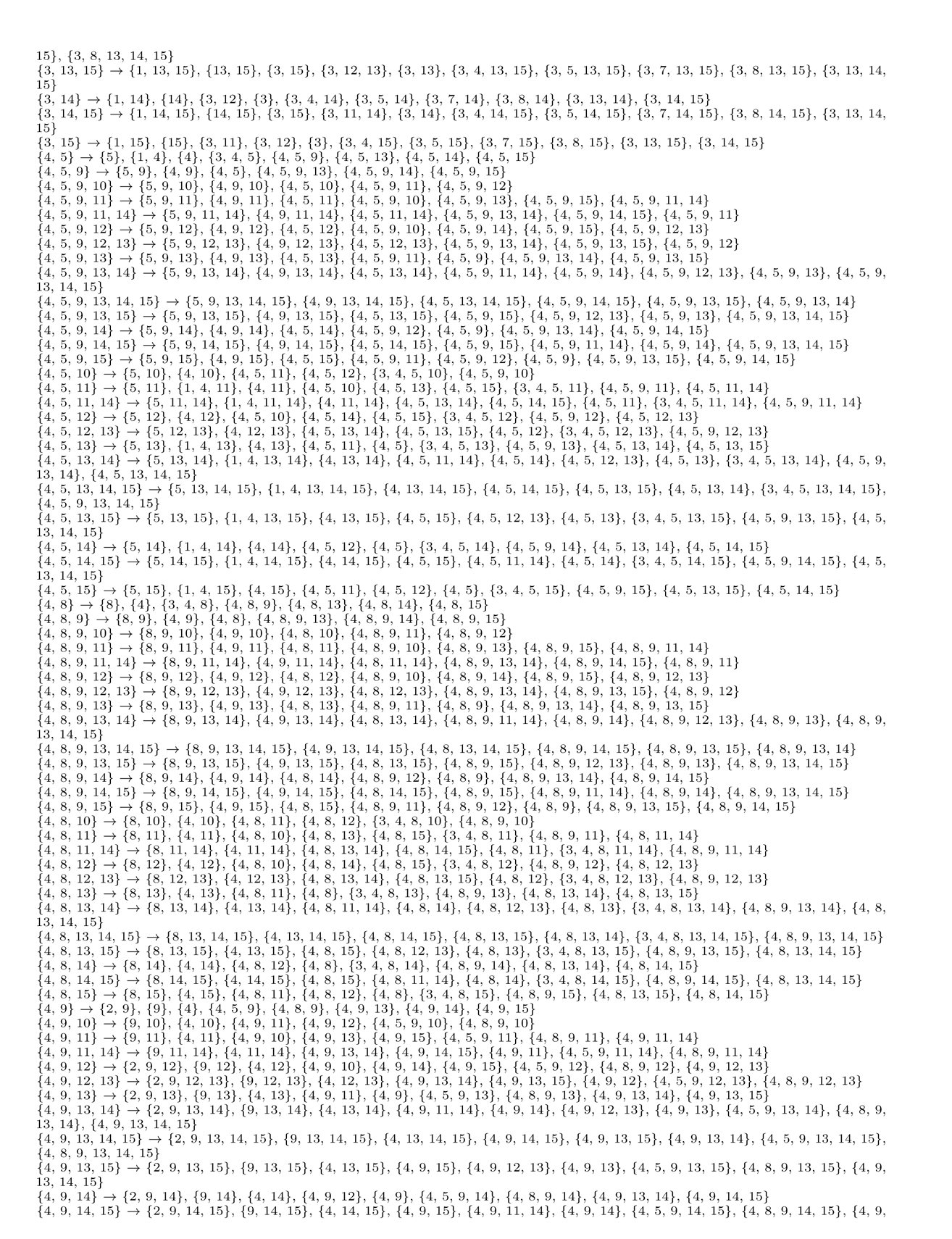

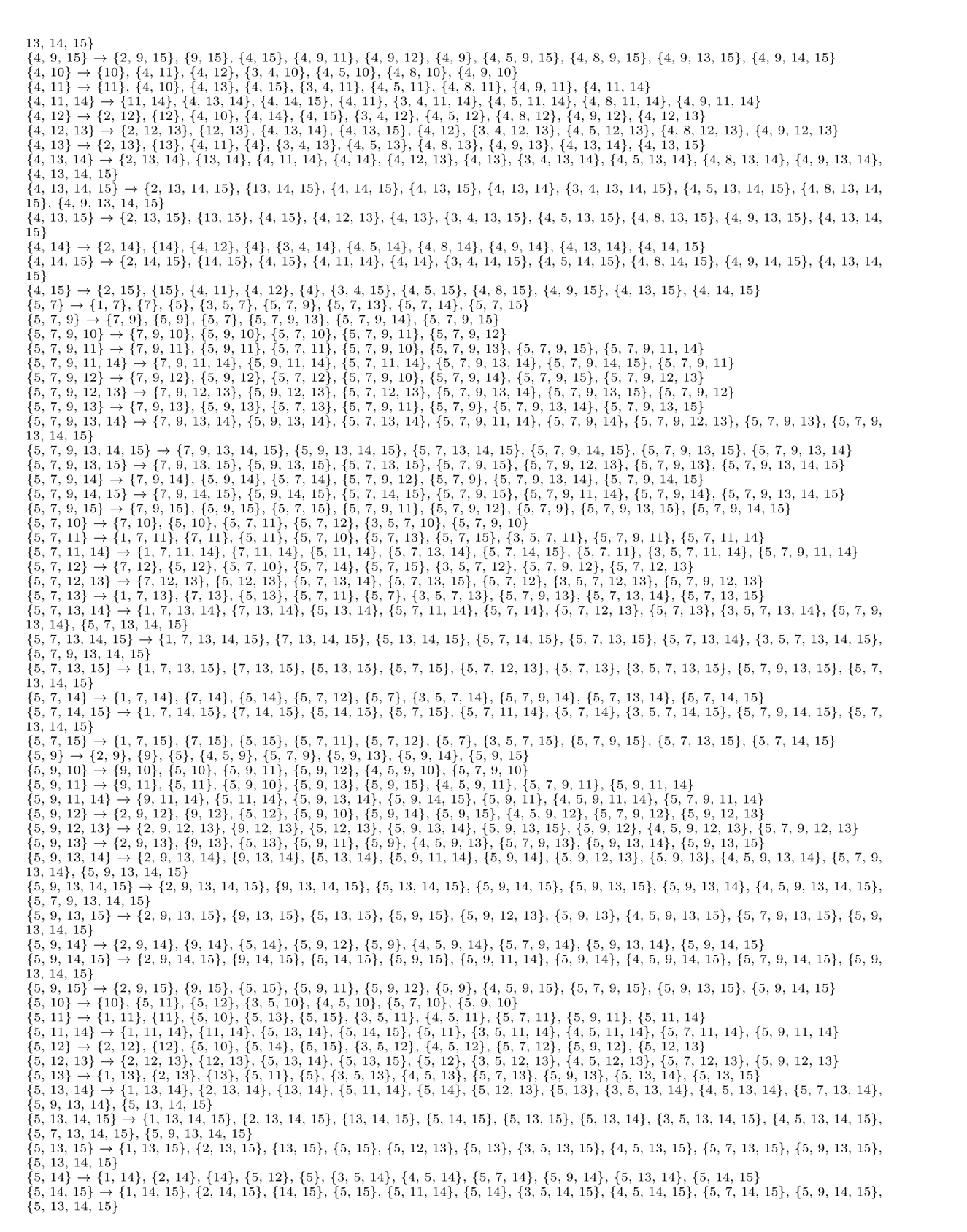

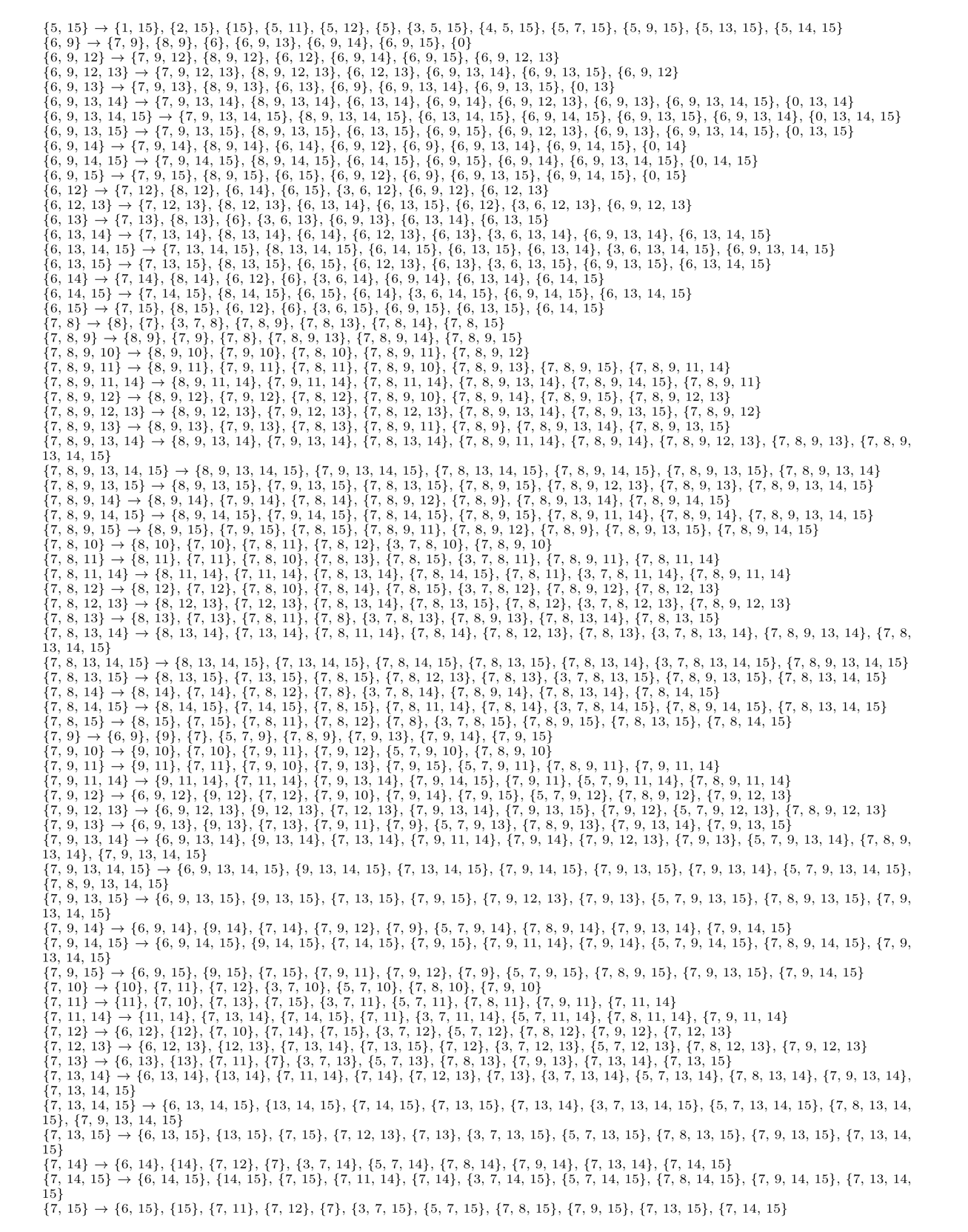

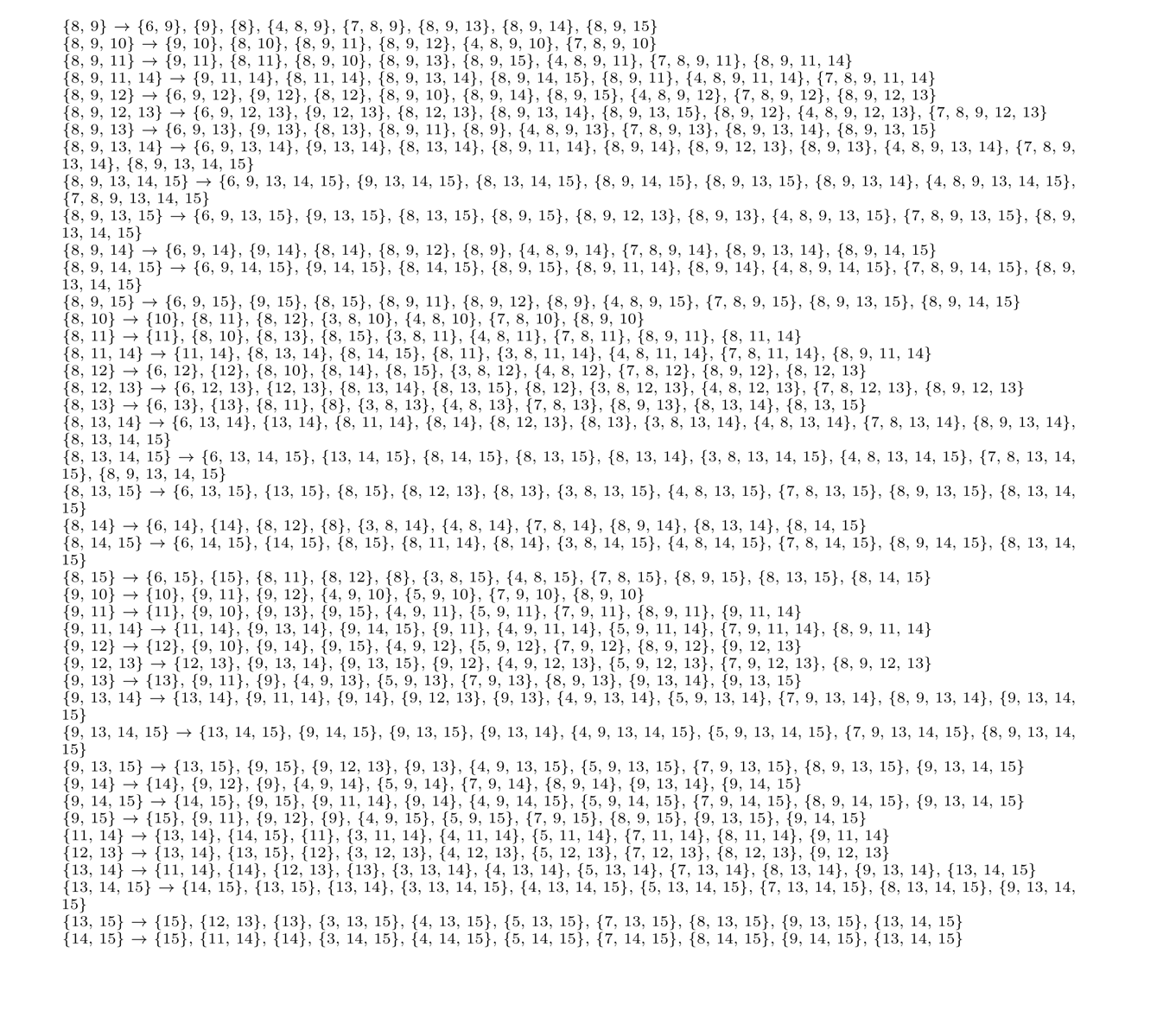

